# Integrating co-expression networks with GWAS to prioritize causal genes in maize

**DOI:** 10.1101/221655

**Authors:** Robert J. Schaefer, Jean-Michel Michno, Joseph Jeffers, Owen Hoekenga, Brian Dilkes, Ivan Baxter, Chad L. Myers

## Abstract

Genome-wide association studies (GWAS) have identified loci linked to hundreds of traits in many different species. Yet, because linkage equilibrium implicates a broad region surrounding each identified locus, the causal genes often remain unknown. This problem is especially pronounced in non-human, non-model species where functional annotations are sparse and there is frequently little information available for prioritizing candidate genes. We developed a computational approach, Camoco, that integrates loci identified by GWAS with functional information derived from gene co-expression networks. Using Camoco, we prioritized candidate genes from a large-scale GWAS examining the accumulation of 17 different elements in maize seeds. Strikingly, we observed a strong dependence in the performance of our approach on the type of coexpression network used: expression variation across genetically diverse individuals in a relevant tissue context (in our case, roots which are the primary elemental uptake and delivery system) outperformed other alternative networks. Two candidate genes identified by our approach were validated using mutants. Our study demonstrates that co-expression networks provide a powerful basis for prioritizing candidate causal genes from GWAS loci, but suggests that the success of such strategies can highly depend on the gene expression data context. Both the software and the lessons on integrating GWAS data with co-expression networks generalize to species beyond maize.

## Background

Genome-wide association studies (GWAS) are a powerful tool for understanding the genetic basis of trait variation. This approach has been successfully applied to hundreds of important traits in different species, including important yield-relevant traits in crops. Sufficiently powered GWAS often identify tens to hundreds of loci containing hundreds of single-nucleotide polymorphisms (SNPs) associated with a trait of interest (McMullen et al., 2009). In *Zea mays* (maize) alone, GWAS have identified nearly 40 genetic loci for flowering time (Buckler et al., 2009), 89 loci for plant height (Peiffer et al., 2014), 36 loci for leaf length (Tian et al., 2011), 32 loci for resistance to southern leaf blight (Kump et al., 2011), and 26 loci for kernel protein (Cook et al., 2012). Despite an understanding of the overall genetic architecture and the ability to statistically associate many loci with a trait of interest, a major challenge has been the identification of causal genes and the biological interpretation of functional alleles associated with these loci.

Linkage disequilibrium (LD), which powers GWAS, acts as a major hurdle limiting the identification of causal genes. Genetic markers are identified by a GWAS, but often reside outside annotated gene boundaries (Wallace et al., 2014) and can be relatively far from the actual causal polymorphism. Thus, a GWA “hit” can implicate many causal genes at each associated locus. In maize, LD varies between 1 kb to over 1 Mb (Gore et al., 2009), and this range can be even broader in other crop species (Morrell et al., 2005; Caldwell et al., 2006). Moreover, there is increasing evidence that gene regulatory regions play a significant role in functional variation leading to causal variants falling outside annotated gene boundaries (Wray, 2007; Wallace et al., 2014). Several quantitative trait loci (QTLs) composed of non-coding sequences have been previously reported in maize (Clark et al., 2006; Castelletti et al., 2014;Louwers et al., 2009). These challenging factors mean that even when a marker is strongly associated with a trait, many candidate genes are equally plausible until a causal polymorphism is identified.

The issues with narrowing a large set of candidate genes to likely causal genes are exacerbated in crop species, where gene annotation is largely incomplete. For example, in maize, only ~1% of genes have functional annotations based on mutant analyses (Andorf et al., 2016). Thus, even when a list of potential candidate genes can be identified for a particular trait, there are very few sources of information that can help identify genes linked to a phenotype. The interpretation and narrowing of large lists of highly associated SNPs with complex traits are now the bottleneck in developing new mechanistic understanding of how genes influence traits.

One informative and easily measurable source of functional information is gene expression. Surveying gene expression profiles in different contexts, such as throughout 3 tissue development or within different genetic backgrounds, helps establish how a gene’s expression is linked to its biological function, including variation in phenotype. Comparing the similarity of two genes’ expression profiles, or co-expression, quantifies the joint response of the genes to various biological contexts, and highly similar expression profiles can indicate shared regulation and function (Eisen et al., 1998). Analysis of co-expression has been used successfully for identifying functionally related genes, including in several crop species (Schaefer et al., 2014; Mochida et al., 2011; Obayashi et al., 2014; Sarkar et al., 2014; Zheng and Zhao, 2013; Ozaki et al., 2010; Swanson-Wagner et al., 2012; Wen et al., 2018; Michno et al., 2018), and has been used to characterize GWAS results in *Arabidopsis thaliana* (Chan et al., 2011; Corwin et al., 2016; Lee and Lee, 2018; Angelovici et al., 2017).

Because co-expression provides a global measure of functional relationships, it can serve as a powerful means for interpreting GWAS candidate loci. Specifically, we expect that variation in several different genes contributing to the same biological process would be associated with a given phenotype (Wolfe et al., 2005; Rotival and Petretto, 2014). Thus, if genetic variation driving the phenotype captured by GWAS is encoded by co-regulated genes, these datasets will non-randomly overlap. Though not all functional relationships are captured with co-expression relationships (Ritchie et al., 2015), these data still provide a highly informative, and sometimes the only, set of clues about genes that otherwise, have not been studied. This principle has been used successfully with other types of networks, for example, protein-protein interactions (Li et al., 2008), and coexpression has been used as a basis for understanding GWAS in mouse and human (Calabrese et al., 2017; Bunyavanich et al., 2014; Taşan et al., 2014; Shim et al., 2017; Baillie et al., 2018).

We developed a freely available, open-source computational framework called Camoco (**C**o-**a**nalysis of **mo**lecular **co**mponents) designed specifically for integrating GWAS candidate lists with gene co-expression networks to prioritize individual candidate genes. Camoco evaluates candidate SNPs derived from a typical GWAS study, then identifies sets of high-confidence candidate genes with strong co-expression where multiple members of the set are associated with the phenotype of interest.

We applied this approach to maize, one of the most important agricultural crops in the world, yielding 15.1 billion bushels of grain in the United States alone in 2016 (USDA, 2016). We specifically focused on quantitative phenotypes measuring the accumulation of 17 different elements in the maize grain ionome (Al, As, B, Ca, Cd, Fe, K, Mg, Mn, Mo, Na, Ni, Rb, S, Se, Sr, and Zn). Plants must take up all elements except carbon and oxygen from the soil, making the plant ionome a critical component in understanding plant environmental response (Baxter, 2010), grain nutritional quality (Guerinot, 2001), and plant physiology (Baxter et al., 2008).

We evaluated the utility of three different types of co-expression networks for supporting the application of Camoco and demonstrate the efficacy of our approach by simulating GWAS to establish maize-specific SNP-to-gene mapping parameters as well as a robust null model for GWAS-network overlap. This approach does indeed confirm overlap between functional modules captured by co-expression networks and GWAS candidate SNPs for the maize grain ionome. We present high-confidence candidate genes identified for a variety of different ionomic traits, test single gene mutants demonstrating the utility of this approach, and, more generally, highlight lessons about the connection between co-expression and GWAS loci from our study that are likely to generalize to other traits and other species.

## Results

### Camoco: A framework for integrating GWAS results and comparing co-expression networks

We developed a computational framework called Camoco that integrates the outputs of GWAS with co-expression networks to prioritize high-confidence causal genes associated with a phenotype of interest. The rationale for our approach is that genes that function together in a biological process that are identified by GWAS should also have non-random structure in co-expression networks that capture the same biological function. Our approach takes, as input, a list of SNPs associated with a trait of interest and a table of gene expression values and produces, as output, a list of high-priority candidate genes that are near GWAS peaks having evidence of strong co-expression with other genes associated with the trait of interest.

There are three major components of the Camoco framework: a module for SNP-to-gene mapping (Figure 1A), tools for construction and analysis of co-expression networks (Figure 1B), and an “overlap” algorithm that integrates GWAS-derived candidate genes with the co-expression networks to identify high-priority candidate genes with strong co-expression support across multiple GWAS loci (Figure 1C) (see Methods for details on each component).

**Figure 1.**
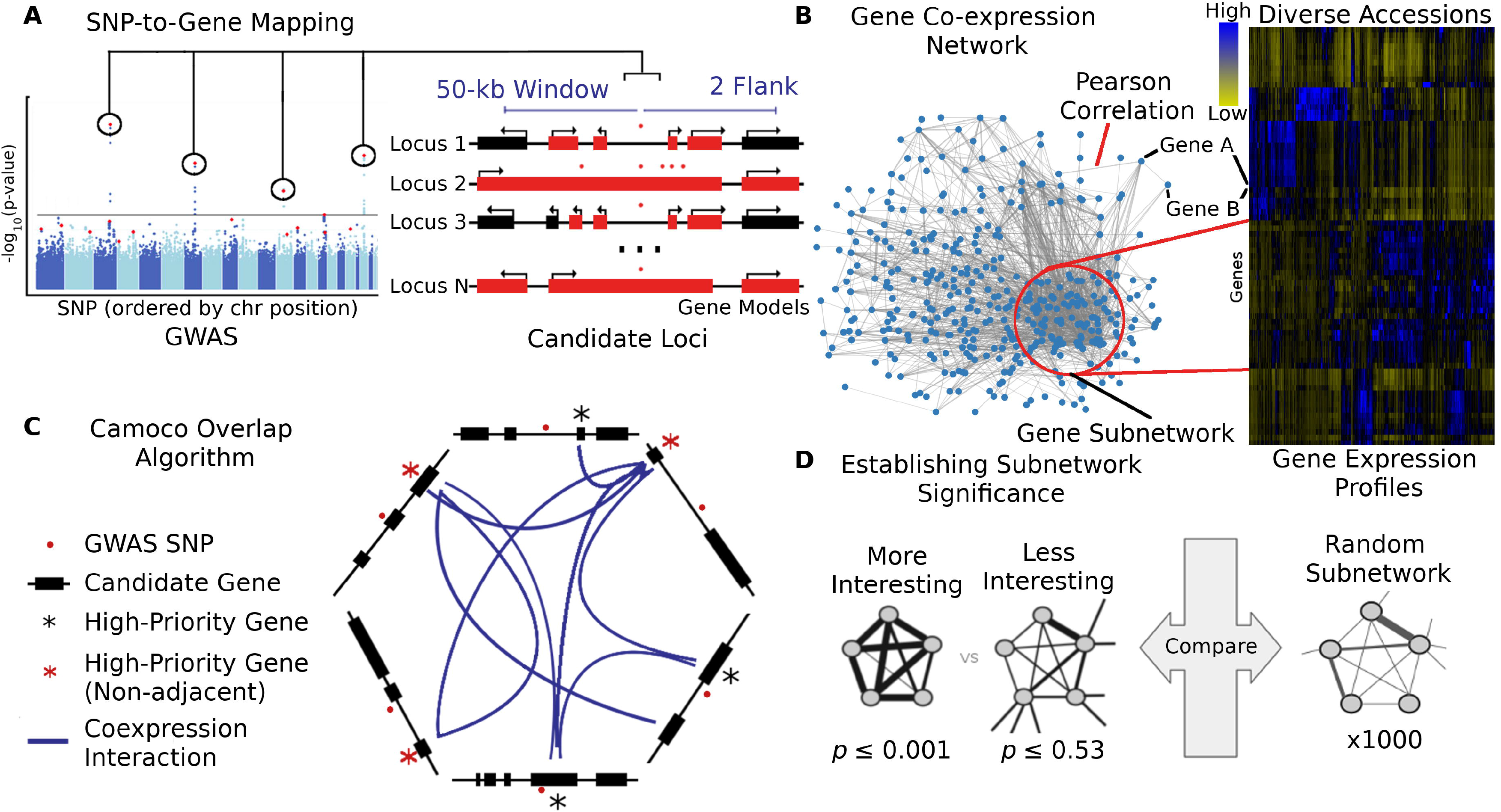
Schematic of the Camoco framework. The Camoco framework integrates genes identified by SNPs associated with complex traits with functional information inferred from co-expression networks. **(A)** A typical GWAS result for a complex trait identifies several SNPs (circled) passing the threshold for genome-wide significance indicating a multigenic trait. SNP-to-gene mapping windows identify a varying number of candidate genes for each SNP. Candidate genes are identified based on user-specified window size and a maximum number of flanking genes surrounding a SNP (e.g., 50-kb and two flanking genes, designated in red). **(B)** Independently, gene co-expression networks identify interactions between genes uncovering an unbiased survey of putative biological co-function. Network interactions are identified by comparing gene expression profiles across a diverse set of accessions (e.g., experimental conditions, tissue, samples). Gene subnetworks indicate sets of genes with strongly correlated gene expression profiles. **(C)** Co-analysis of co-expression interactions among GWAS trait candidate genes identifies a small subset of genes with strong network connections. Blue lines designate genes that have similar co-expression patterns indicating co-regulation or shared function. Starred genes are potential candidate genes associated with GWAS traits based on SNP-to-gene mapping and co-expression evidence. Red stars indicate genes that are not the closest to the GWAS SNP (non-adjacent) that may have been missed without co-expression evidence. **(D)** Statistical significance of subnetwork interactions is assessed by comparing co-expression strength among genes identified from GWAS datasets to those from random networks containing the same number of genes. In the illustrated case, the more interesting subnetwork has both high density as well as locality.

The overlap algorithm uses two network scoring metrics: subnetwork density and subnetwork locality. Subnetwork density measures the average interaction strength between all pairwise combinations (i.e. unthresholded) of genes near GWAS peaks. Specifically, density is obtained by computing the mean of raw interaction scores among all pairs of genes in the subnetwork and normalizing by the subnetwork size (Eq. 1). Subnetwork locality measures the proportion of significant (Z≥3) co-expression interactions among genes within a GWAS-derived subnetwork (local interactions) as compared to the number of global interactions with other genes in the genome (global interactions). Specifically, locality is obtained by first fitting a linear regression between all genes’ local degree (among the subnetwork of interest) and their global degree and measuring the mean of the residual for genes in the subnetwork (Eq. 2). Density and locality metrics can be calculated on whole subnetworks or on a gene-specific basis to prioritize candidate genes by factoring out each gene’s contribution to the subnetwork (Eq. 3 and Eq. 4) (see Methods for details). For a given input GWAS trait and coexpression network, the statistical significance for both density and locality is determined by generating a null distribution based on randomly generated GWAS traits (*n* = 1,000) with the same number of implicated loci and corresponding candidate genes. The resulting null distribution is then used to derive a *p*-value for the observed subnetwork density and locality for all putative causal genes (Figure 1D). Thus, for a given input GWAS trait, Camoco produces a ranked list of candidate causal genes for both network metrics and a corresponding false discovery rate (FDR) that indicates the significance of the observed overlap between each candidate causal gene’s co-expression network neighbors and the set of genes under implicated loci. Using this integrated approach, the number of candidate genes prioritized for follow-up validation is reduced substantially relative to the initial set of genes under implicated loci.

Camoco allows users to build, validate, and analyze datasets using common file-types for gene-expression, GWAS and species-specific reference data (e.g. OBO, FASTA, GFF). Our tool formalizes the integration of GWAS data with co-expression networks by offering systematic SNP-to-gene mapping parameters, which can be evaluated using simulated GWAS gold standard datasets. Camoco also corrects for artifacts (such as cis-co-expression bias) that arise from integrating GWAS and co-expression data. The framework offers a unified command line interface to the components described above but can also be used through its python API to integrate into other workflows. Our method can be applied to any trait and species for which GWAS has been completed and sufficient gene expression data exist to construct a co-expression network.

#### Generating co-expression networks from diverse transcriptional data

A co-expression network that is derived from the biological context generating the phenotypic variation subjected to GWAS is a key component of our approach. A well-matched co-expression network will describe the most relevant functional relationships and identify coherent subsets of GWAS-implicated genes. We and others have previously shown that co-expression networks generated from expression data derived from different contexts capture different functional information (Schaefer et al., 2014; Swanson-Wagner et al., 2012). For example, experiments measuring changes in gene expression can explore environmental adaptation, developmental and organ-based variation, or variation in expression that arises from population and ecological dynamics (see (Schaefer et al., 2016) for review). For some species, published data contain enough experimental accessions to build networks from these different types of expression experiments (the term accession is used here to differentiate samples, tissues, conditions, etc.). We reasoned that these different sources of expression profiles likely have a strong impact on the utility of the co-expression network for interpreting genetic variation captured by GWAS. Using this rationale, we constructed several different co-expression networks independently and assessed the ability of each to produce high-confidence discoveries using our Camoco framework.

Three co-expression networks representing three different biological contexts were built. The first dataset targeted expression variation that exists between diverse maize accessions built from whole-seedling transcriptomes on a panel of 503 diverse inbred lines from a previously published dataset characterizing the maize pan-genome (Hirsch et al., 2014) (called the ZmPAN network hereafter). Briefly, Hirsch et al. chose these lines to represent major heterotic groups within the United States, sweet corn, popcorn, and exotic maize lines and measured gene expression profiles for seedling tissue as a representative tissue for all lines. The second dataset examined gene expression variation from a previous study characterizing different tissues and developmental time points (Stelpflug et al., 2015). Whole-genome RNA-Seq transcriptome profiles from 76 different tissues and developmental time points from the maize reference accession B73 were used to build a network representing a single-accession expression map (called the ZmSAM network hereafter). Finally, we created a third dataset as part of the ionomics GWAS research program. These data measure gene expression variation in the root, which serves as the primary uptake and delivery system for all the measured elements (Chao et al., 2011; Baxter, 2010; Baxter and Dilkes, 2012). Gene expression was measured from mature roots in a collection of 46 genotypically diverse maize inbreds (called the ZmRoot network hereafter). All datasets used here were generated from whole-genome RNA-Seq analysis, although Camoco could also be applied to microarray-derived expression data.

Co-expression networks for each dataset were constructed from gene expression matrices using Camoco (see Methods for specific details on building these networks). Once built, several summary statistics were evaluated from interactions that arise between genes in the network (Supp. Figure 1–3). Co-expression was measured among genes within the same Gene Ontology (GO) term to establish how well density and locality captured terms with annotated biological functions (Table 1; Supp. Table 1). Indeed, we observed enrichment for a large number of GO terms for both metrics in all three networks as well similar levels of enriched modules derived from a graph clustering approach (Table 2; Supp. Table 2;), supporting their ability to capture functionally related genes (See Discussion, Supplementary Text and Supp. Table 3). Indeed, we observed enrichment for a large number of GO terms for both metrics in all three networks as well similar levels of enriched modules derived from a graph clustering approach (Table 2; Supp. Table 2;), supporting their ability to capture functionally related genes (See Discussion, Supplementary Text and Supp. Table 3).

**Table 1.** **Significantly co-expressed GO terms.** Co-expression was measured among genes within each GO term that had Co-expression data in each network using both density (Eq. 1) and locality (Eq. 2). Significance of Co-expression metrics was assessed by comparing values to 1,000 random gene sets of the same size.

| | Number Significant ($p \leq 0.01$) GO Terms (n = 1078) | | | |
| --- | --- | --- | --- | --- |
|  | Density | Locality | Both Scores | Either Score |
| ZmPAN | 451 (41%) | 539 (50%) | 312 (29%) | 678 (63%) |
| ZmSAM | 365 (34%) | 437 (40%) | 234 (21%) | 568 (53%) |
| ZmRoot | 573 (53%) | 331 (31%) | 278 (26%) | 626 (58%) |

**Table 2.** **Gene Co-expression network cluster assignments.** Gene clusters were calculated by running the Markov Cluster (MCL) algorithm on the Co-expression matrix. Cluster values designate network specific gene clusters and are not compared across networks.

|  | Network Clusters |  |  |
| --- | --- | --- | --- |
| | Num Cluster: ($10 \geq n > 100$) | Num Clusters: ($n \geq 100$) | Num Clusters ($n \geq 10$) Enriched for GO Terms ($p \leq 0.01$) |
| ZmPAN | 76 | 18 | 71 |
| ZmSAM | 160 | 10 | 115 |
| ZmRoot | 150 | 10 | 106 |

#### Accounting for *cis* gene interactions

Camoco integrates GWAS candidates with co-expression interactions by directly assessing the density or locality of interactions among candidate genes near GWAS SNPs. However, the process of mapping SNPs to surrounding candidate genes has inherent complications that can strongly influence subnetwork co-expression calculations. While we assume that the majority of informative interactions among candidate genes are between GWAS loci, *cis*-regulatory elements and other factors can lead to co-expression between linked genes and produce skewed distributions in density and locality calculations, which can in turn bias co-expression statistics. Identifying significant overlap between GWAS loci and co-expression networks requires a distinction between co-expression among genes that are in close proximity to one another on a chromosome (*cis*) compared to those genes that are not (*trans*).

To assess the impact of *cis* co-expression, network interactions for genes located on different chromosome (*trans* interactions) were compared to *cis* interactions for pairs of genes less than 50 kb apart. The distributions of the two groups indicate that *cis* genes are more likely to have a strong co-expression interaction score than *trans* genes (Figure 2). This bias toward *cis* genes is especially pronounced for strong positive co-expression, where we observed substantially stronger enrichment for linked gene pairs compared to *trans* genes (e.g., z-score ≥ 3; see Figure 2 inset).

**Figure 2.**
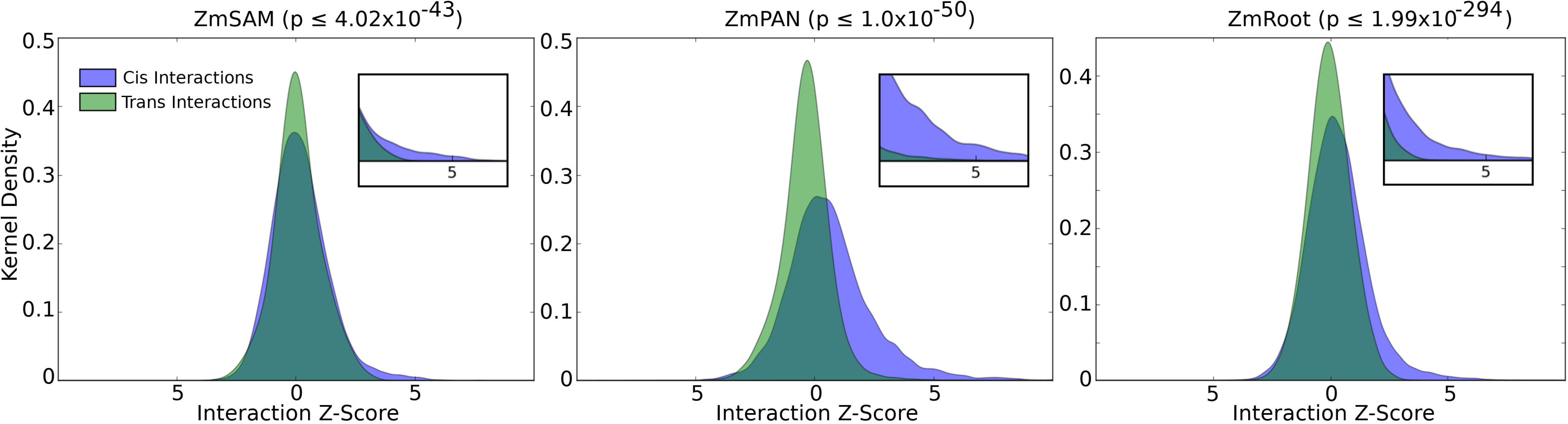
*Cis* vs. *trans* co-expression network interactions. Comparing distributions of co-expression network interaction scores between *cis* and *trans* sets of genes. Distribution densities of *trans* gene pairs (green) show interactions between genes on separate chromosomes. Distribution densities of *cis* gene pairs (blue) show interactions between genes with less than 50 kb intergenic distance. Inset figures show z-score values greater than 3. Non-parametric *p*-values were calculated between co-expression values taken from *cis* and *trans* distributions (Mann-Whitney U test).

The enrichment of significant co-expression among *cis* genes, likely due to shared *cis*-regulatory sequences or closely encoded clusters of functionally related genes, prompted us to remove *cis* interactions when examining co-expression relationships among candidate genes identified by GWAS SNPs in Camoco. To account for the bias of strong co-expression among *cis* genes, only interactions among pairs of genes originating from unlinked SNPs (i.e. *trans*) were included in density and locality calculations when evaluating GWAS results (see Methods).

#### Evaluation of the Camoco framework

To explore the limits of our approach, we examined factors that influence overlap detection between co-expression networks and genes linked to GWAS loci. In an idealized scenario, SNPs identified by GWAS map directly to true causal genes, all of which exhibit strong co-expression network interactions with each other (Figure 3). But in practice, SNPs can affect regulatory sequences or be in linkage disequilibrium (LD) with the functionally important allele, leading to a large proportion of SNPs occurring outside of genic regions (Wallace et al., 2014).

**Figure 3.**
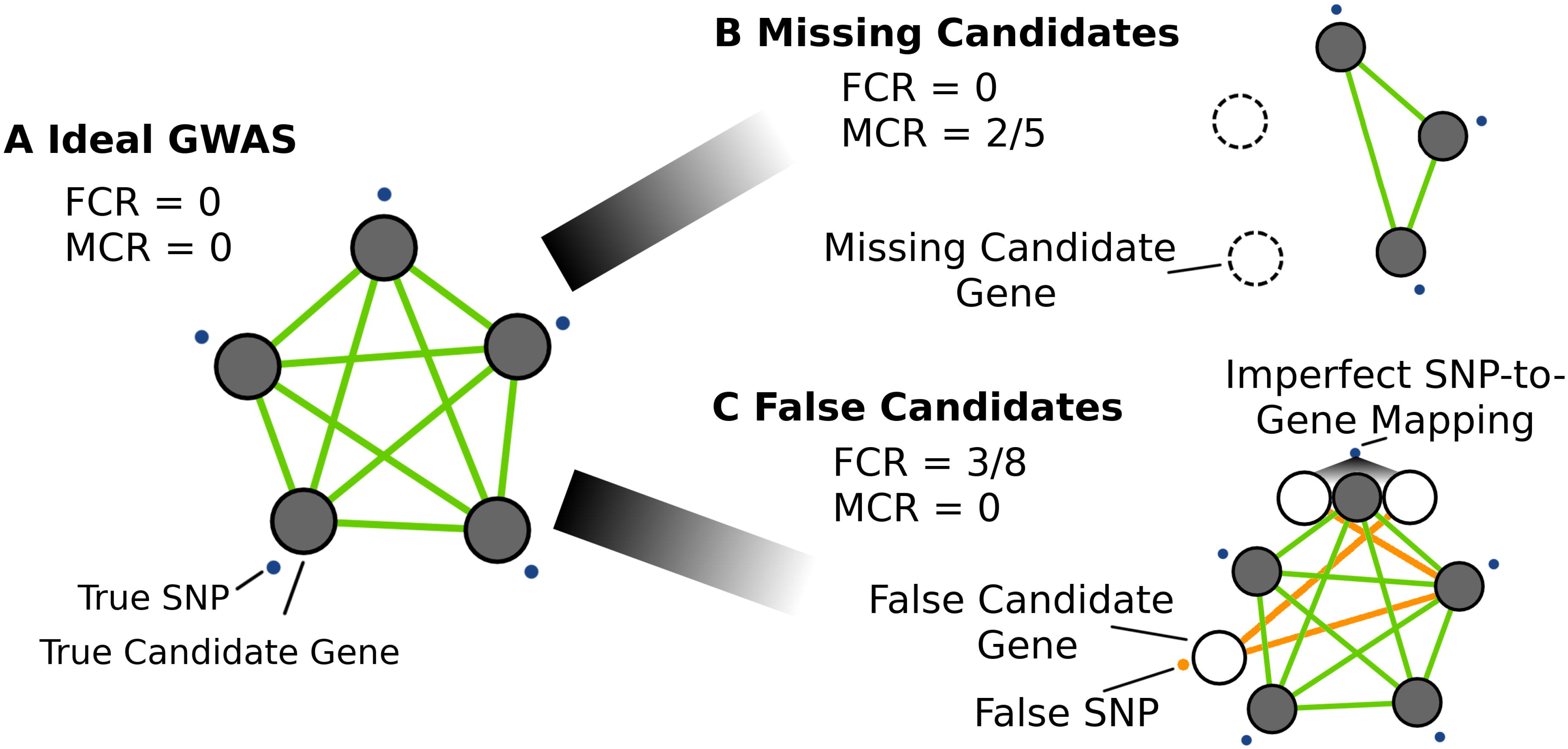
Simulating GWAS-network overlap using GO terms. Several GWAS scenarios were simulated to assess the effect of noise on co-expression network overlap. Panel **(A)** shows an ideal GWAS, where SNPs (blue points) map directly to candidate genes within the same biological process (i.e., a GO term) and have strong co-expression (green lines). Signal is defined as the co-expression among the genes exclusive to the GO term. Noise in the overlap between GWAS and co-expression networks was introduced by varying two parameters: the missing candidate gene rate (MCR) and false candidate gene rate (FCR). Panel **(B)** demonstrates the effect of a large proportion of missing candidate genes (MCR = 2/5) on network signal. Likewise, panel **(C)** shows the effect of false candidate genes (FCR) on network overlap, either through false positive GWAS SNPs (orange points) or through imperfect SNP-to-gene mapping (FCR = 3/8). Orange lines designate the additional candidate genes that introduce co-expression noise that impedes the identification of network structure.

We evaluated two major challenges that influence SNP-to-gene mapping. The first is the total number of functionally related genes in a subnetwork, representing the fraction of genes involved in a biological process that are simultaneously identified by GWAS. In cases where too few genes represent any one of the underlying causal processes, our proposed approach is not likely to perform well—for example, consider the situation when GWAS identifies a single locus in a ten-gene biological process due to incomplete penetrance, limited allelic variation in the mapping population, or extensive gene-by-environment interactions. We refer to this source of noise as the *missing candidate gene rate* (*MCR*) or, in other words, the fraction of genes involved in the causal process not identified by the GWAS in question (Figure 3B; Eq. 5).

The second key challenge in identifying causal genes from GWAS loci is instances where associated SNPs each implicate a large number of non-causal candidate genes. Thus, in cases where the linked regions are large (i.e., imperfect SNP-to-gene mapping), the framework’s ability to confidently identify subnetworks of highly co-expressed causal genes may be compromised. One would expect to find scenarios where the proposed approach does not work simply because there are too many non-causal genes implicated by linkage within each GWAS locus, such that the co-expression signal among the true causal genes is diminished by the false candidates linked to those regions. We refer to this source of noise as the *false candidate gene rate* (*FCR*), the fraction of all genes linked to GWAS-implicated loci that are not causal genes (Figure 3C; Eq. 6).

To explore the limits of our co-expression-based approach with respect to these factors, we simulated scenarios where we could precisely control both MCR and FCR. In practice, neither of these quantities can be controlled; MCR is a function of the genetic architecture of the phenotype as well as the degree of power within the study population of interest, and FCR is a function of recombination frequency in the GWAS population.

We evaluated the expected performance of the Camoco framework for a range of each of these parameters by simulating ideal GWAS scenarios using GO terms with significantly co-expressed genes (*p ≤* 0.05; Table 1). These ideal cases were then subjected to processes where either a subset of genes was replaced by random genes (i.e., to simulate MCR but conserve term size) or additional functionally unrelated genes were added using SNP-to-gene mapping (i.e., to simulate FCR introduced by linkage). In both cases, simulated GWAS candidates (i.e. genes annotated to our selected GO terms) were subjected to varying levels of either FCR or MCR while tracking the number of GO terms that remained significantly co-expressed at each level. These simulations enabled us to explore a broad range of settings for these key parameters and establish whether our proposed approach had the potential to be applied in maize.

#### Simulated GWAS datasets show robust co-expression signal to MCR and FCR

Subnetwork density and locality were measured for GO terms with significantly co-expressed genes containing between 50 and 150 genes in each network at varying levels of MCR (see Supp. Table 4). At each MCR level, density and locality among the remaining genes were compared to 1,000 random sets of genes of the same size. The proportion of initial GO terms that remained significantly co-expressed was recorded for each network (see Figure 4, red curve; see Supp. Figure 4A for absolute term numbers). GO terms were also split into two starting groups based on strength of initial co-expression: moderate (0.001 < *p* ≤ 0.05; blue curve) and strong (*p* ≤ 0.001; violet curve).

**Figure 4.**
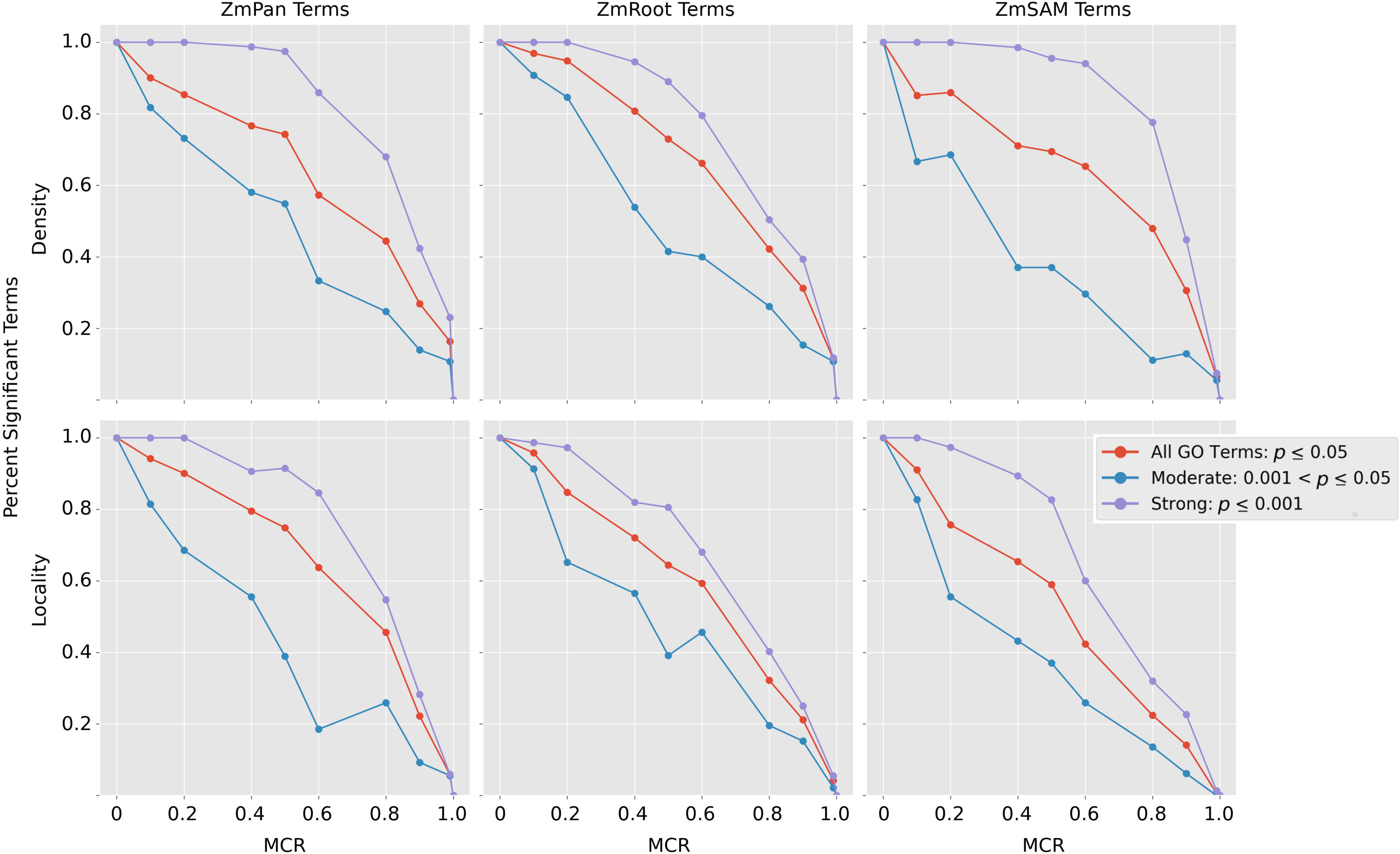
Strength of co-expression among GO terms at varying levels of MCR. Subnetwork density and locality were measured for all GO terms with strong initial co-expression (*p* ≤ 0.05) comparing co-expression in GO terms to 1,000 random networks of the same size. Co-expression density and locality were then compared again (*n* = 1,000) with varying missing candidate rate (MCR), where a percentage of genes was removed from the term and replaced with random genes to conserve GO term size. Curves decline with increased MCR as the proportion of GO terms with significantly co-expressed genes (*p* ≤ 0.05, *n* = 1,000) decreases compared to the initial number of strongly co-expressed terms in each network (red curve). GO terms in each network were also split into two subsets based on initial co-expression strength: “strong,” (initial co-expression *p* ≤ 0.001; blue curve), and “moderate,” (initial co-expression 0.001 < *p* ≤ 0.05; violet curve).

As expected, strength of co-expression among GO terms decreased as MCR increased. Figure 4 shows the decay in the proportion of GO terms that exhibit significant co-expression at increasing levels of MCR (red curve). In general, the decay of signal is similar between density and locality, where signal initially decays slowly until approximately 60% MCR, when signal quickly diminishes.

In all three networks, GO terms with stronger initial co-expression were more robust to MCR. Signal among GO terms with strongly co-expressed genes (*p* ≤ 0.001; violet curve) decayed at a substantially lower rate than GO terms with a more moderate signal, indicating that this approach is robust for GWAS datasets with moderate levels of missing genes when co-expression among true candidate genes is strong. Co-expression signal in relation to MCR was also compared between GO terms split by the number of genes within the term (see Supp. Figure 4B–C), which did not influence the rate at which co-expression signal decayed.

Likewise, the effect of FCR was simulated. GO terms with between 50 and 150 genes (MCR = 0) with significant co-expression among member genes (*p* ≤ 0.05; see Supp. Table 4) were selected. The nucleotide position of the starting base pair of each true GO term gene was used as input for our SNP-to-gene mapping protocol for identifying GWAS candidates (see Methods). Subnetwork density and locality were calculated for the simulated candidate genes corresponding to each SNP-to-gene mapping combination, in each network, to evaluate the decay of co-expression signal as FCR increases (Figure 5).

**Figure 5.**
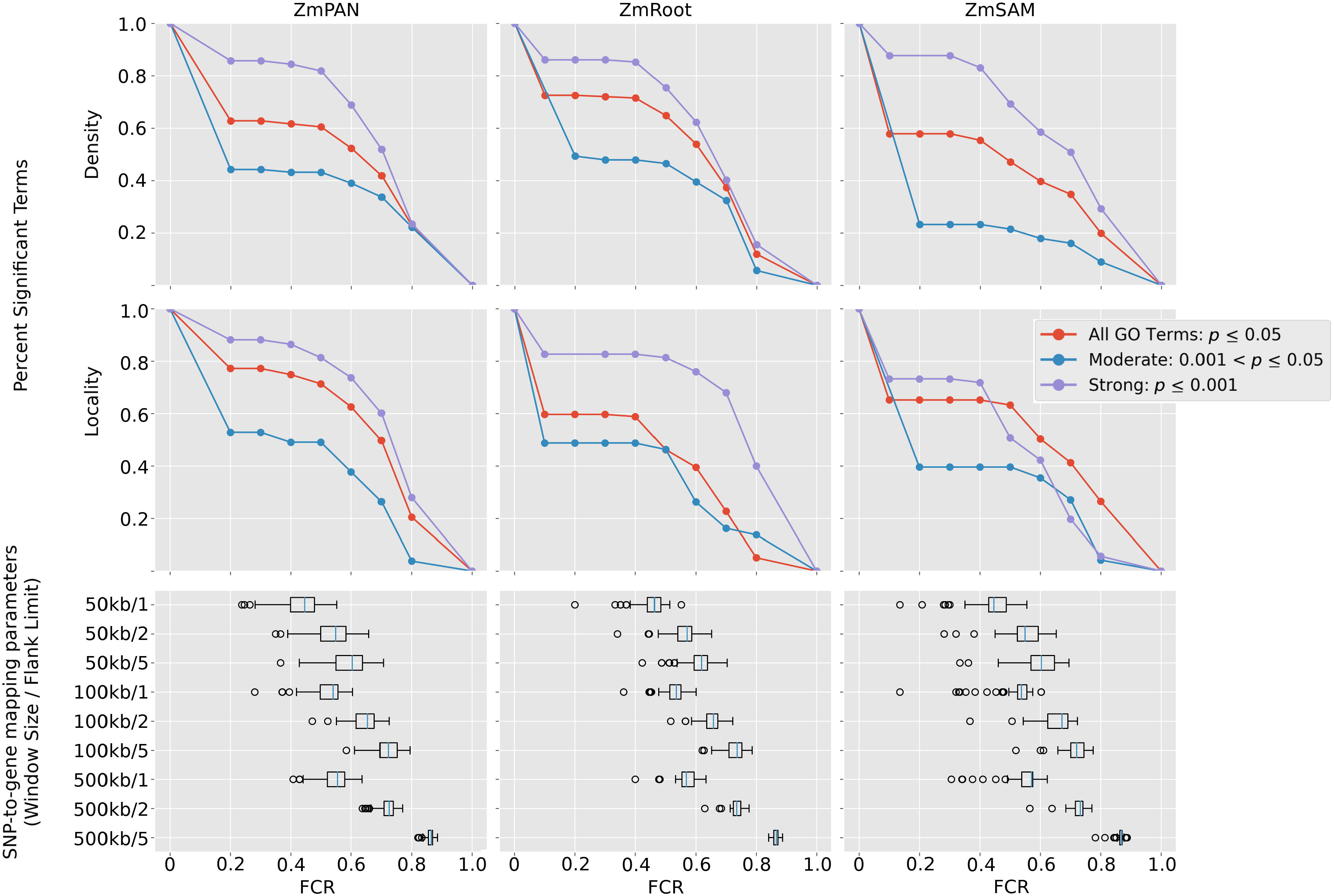
Strength of co-expression among GO terms at varying levels of FCR. GO terms with significantly co-expressed genes (density or locality *p*-value ≤ 0.05) were used to simulate the effect of FCR on GWAS results. False candidates were added to GO terms by including flanking genes near true GO term genes according to SNP-to-gene mapping (window) parameters. Box plots show effective FCR of GO terms at each SNP-to-gene mapping parameter. Signal plots show the proportional number of GO terms that remain significant at FCR ≥ *x* (red curve). GO terms in each network were also split into two subsets based on initial co-expression strength: “strong,” (initial co-expression *p* ≤ 0.001; blue curve), and “moderate,” (initial co-expression 0.001 < *p* ≤ 0.05; violet curve).

Candidate genes were added by varying the window size for each SNP up to 50 kb, 100 kb, and 500 kb upstream and downstream and by varying the maximum number of flanking genes on each side to one, two, and five. Given the number of additional candidate genes introduced at each SNP-to-gene mapping combination, FCR was calculated for each GO term at each window size (see Figure 5 box plots).

Co-expression signal in relation to FCR was assessed by comparing subnetwork density and locality for each GO term at different SNP-to-gene mapping parameters for each of the three co-expression networks to random subnetworks with the same number of genes (*n* = 1,000) (Figure 5, top). The proportion of GO terms with significantly co-expressed genes decayed at higher levels of FCR (see Supp. Figure 5A for absolute term numbers). The minimum FCR level ranged from 1% to 80% across all GO terms, but for most GO terms was ~50% as the most stringent SNP-to-gene mapping (50 kb/one flank) approximately doubled the number of candidate genes. Two additional scenarios were considered in which signal was further split based on the initial co-expression strength: “moderate” (0.001 < *p* < 0.05; blue curve) and “strong” (*p* ≤ 0.001; violet curve).

Despite high initial false candidate rates, co-expression signal among GO terms remained significant even at 60–70% FCR. Similar to the results with MCR, GO terms with stronger initial co-expression were more likely to remain significantly co-expressed at higher FCR levels. Co-expression signal in relation to FCR was also compared between GO terms split by the number of genes in the term (see Supp. Figure 5B–C), which did not differentiate the rate at which co-expression signal decayed.

In cases where true candidate genes identified by GWAS were strongly co-expressed, as simulated here, a substantial number of false positive SNPs or an introduction of false candidate genes through uncertainty in SNP-to-gene mapping, can be tolerated, and network metrics still detected the underlying co-expressed gene sets using our method. These results indicate that in GWAS scenarios where the majority of SNPs do not perfectly resolve to candidate genes, systematic integration with co-expression networks can efficiently filter out false candidates introduced by SNP-to-gene mapping if the underlying causal loci are linked to genes that are strongly co-expressed with each other. Moreover, in instances where several intervening genes exist between strongly associated SNPs in LD with each other and the true causative allele, true causal candidates can be detected using co-expression networks as a functional filter for candidate gene identification.

The potential for using this approach, however, is highly dependent on the LD of the organism in question, the genetic architecture of the trait being studied, and the degree of co-expression between causative loci. Simulations provide insight into the feasibility of using Camoco to evaluate overlap between co-expression networks and GWAS as well as a survey of the SNP-to-gene mapping parameters that should be used when using this approach (see Discussion for more details). In the context of maize, simulations performed here suggest that systematic integration of co-expression networks to interpret GWAS results will increase the precision with which causal genes associated with quantitative traits in true GWAS scenarios can be identified.

#### High-priority candidate causal genes under ionomic GWAS loci

Identifying the biological processes underlying the elemental composition of plant tissues, also known as the ionome, can lead to a better understanding of plant adaptation as well as improved crops (Baxter and Dilkes, 2012). High-throughput analytic approaches such as inductively coupled plasma mass spectrometry (ICP-MS) are capable of measuring elemental concentrations for multiple elements and are scalable to thousands of accessions per week. Using ICP-MS, we analyzed the accumulation of 17 elements in maize kernels described in depth by Ziegler et al. (Ziegler et al., 2017). Briefly, kernels from the nested association mapping (NAM) population were grown in four geographic locations (McMullen et al., 2009). To reduce environmental-specific factors, the SNPs used in this study were from the GWAS performed on the all-location models. Approximately 30 million SNPs and small copy-number variants were projected onto the association panel and used to perform a GWAS for each of the 17 elements. SNPs were tested for significance of association for each trait using resampling model inclusion probability (Valdar et al., 2009) (RMIP ≤ 0.05; see Methods). For each element (trait), significantly associated SNPs were used as input to Camoco to generate candidate genes from the maize filtered gene set (FGS; *n* = 39,656) using a range of SNP-to-gene mapping parameters: 50-kb, 100-kb, and 500-kb windows (up/downstream) limited each to one, two, or five flanking genes (up/downstream of SNP; see Figure 1A). In total, 4,243 statistically significant SNPs were associated with maize grain ionome traits. Summing the potential candidate genes across all 17 traits implicates between 5,272 and 22,927 unique genes depending on the SNP-to-gene mapping parameters used (between 13% and 57% of the maize FGS, respectively; See Supp. Table 5). On average, each trait’s significantly associated SNPs identified 119 non-overlapping windows across the ten chromosomes of maize (i.e., effective loci; see Methods), and these implicate an average of 613 candidate genes per element (Methods).

Given the large number of candidate genes associated with elemental accumulation, we used Camoco to integrate network co-expression with effective loci identified by GWAS for each of the 17 elemental traits separately. By combining candidate gene lists with the three gene expression datasets (ZmPAN, ZmRoot, and ZmSAM) and two co-expression network metrics (locality and density), high-priority candidate genes driving elemental accumulation in maize were identified (see Figure 1C). For each network-trait combination, Camoco identified a ranked list of prioritized candidate causal genes, each associated with an FDR that reflects the significance of co-expression connecting that candidate gene to genes near other loci associated with the same trait (Supp. Table 6). We defined a set of high-confidence discoveries by reporting candidates that were discovered at a FDR ≤ 30% in at least two SNP-to-gene mapping parameter settings (e.g., 50 kb/one flank and 100 kb/one flank), denoted as the high-priority overlap (HPO) set (see Supp. Table 7 and Methods).

By these criteria, we found strong evidence of co-expression for 610 HPO genes that were positional candidates across the 17 ionomic traits measured (1.5% maize FGS). The number of HPO genes discovered varied significantly across the traits we examined, with between 2 and 209 HPO genes for a given element considering either density or locality in any network (Figure 6; Either:Any column). HPO genes discovered by Camoco were often non-adjacent to GWAS effective loci, either having genes intervening between the HPO candidate or that were closer to the GWAS-implicated locus (Figure 1C), demonstrating that Camoco often identifies candidates with strong co-expression evidence that would not have been selected by choosing the closest positional candidate.

**Figure 6.**
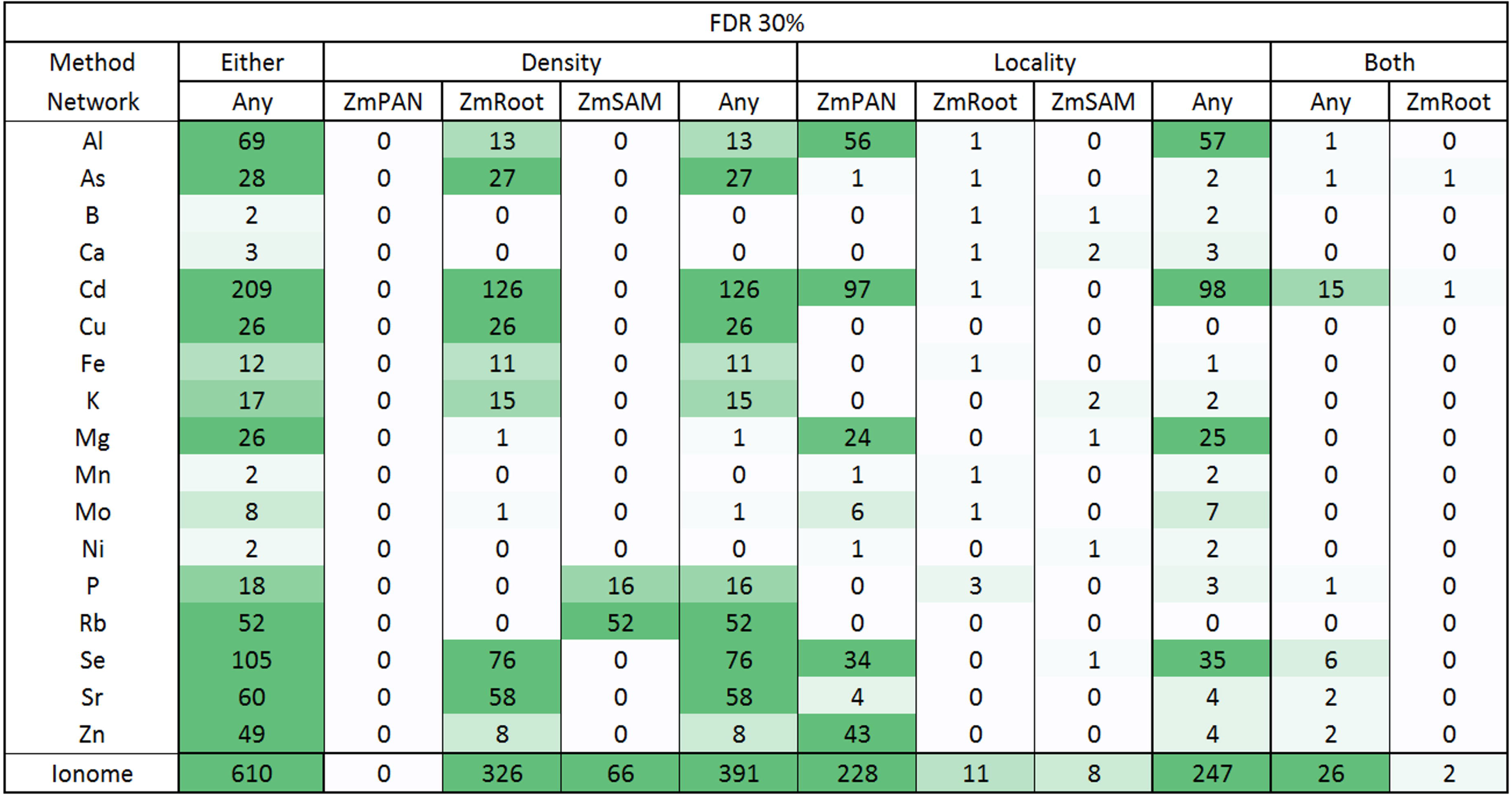
Maize grain ionome high-priority candidate genes heatmap summary. Gene-specific density and locality metrics were compared to (*n* = 1,000) random sets of genes of the same size to establish a 30% FDR. Genes were considered candidates if they were observed at two or more SNP-to-gene mappings (i.e., HPO). Candidates in the “Either” column are HPO genes discovered by either density or locality in any network. The number of genes discovered for each element is further broken down by co-expression method (density, locality, both) and by network (ZmPAN, ZmSAM, ZmRoot). Candidates in the “Both” column were discovered by density and locality in the same network or in different networks (Any). Note: zero elements had HPO genes using “Both” methods in the ZmPAN and ZmSAM networks.

### Genotypically diverse networks support stronger candidate gene discoveries than tissue atlases

The variation in the number of genes discovered by Camoco depended on which co-expression network was used as the basis for discovery. The ZmRoot co-expression network proved to be the strongest input, discovering genes for 15 of the 17 elements (absent in Ni and Rb) for a total of 335 HPO genes, ranging from 1 to 126 per trait (Supp. Table 7). In contrast, the ZmSAM network, which was constructed based on a tissue and developmental expression atlas collected exclusively from the B73 accession, supported the discovery of candidate genes for only 8 elements (B, Ca, K, Mg, Ni, P, Rb, and Se) for a total of 74 HPO genes, ranging from 1 to 52 per trait (Supp. Table 7). The ZmPAN network, which was constructed from whole seedlings (pooled tissue) across 503 different accessions, provided intermediate results, supporting high-confidence candidate discoveries for 10 elements (Al, As, Cd, Mg, Mn, Mo, Ni, Se, Sr, and Zn) for a total of 228 HPO genes, ranging from 1 to 97 per trait (Supp. Table 7). The relative strength of the different networks for discovering candidate causal genes was consistent even at stricter FDR thresholds (e.g., FDR ≤ 0.10; Supp. Table 7).

#### Network co-expression metrics provide complementary information and most candidate causal genes are trait specific

Both density and locality were assessed on a gene-specific level to measure the strength of a given candidate causal gene’s co-expression relationships with genes in other GWAS-identified loci (see Eq. 3 and Eq. 4) (see Figure 6, Density:Any and Locality:Any). Interestingly, the high-confidence genes identified by the two approaches were largely complementary, in terms of both which traits and which network they produced results for. Indeed, when we measured the direct correlation of gene-specific density and locality measures across several GWAS traits and GO terms, we observed very weak positive but significant correlations (Supp. Figure 6). We observed that the utility of the locality metric appeared to be associated with the number of accessions used to construct the network (Supp. Table 8, see Discussion). One important question is the extent to which putative causal genes overlap across different ionomic traits. It is plausible that some mechanisms affecting elemental accumulation are shared by multiple elements. However, most of the discovered HPO genes are element specific, with relatively little overlap between elements (Supp. Figure 7; Supp. Table 9).

### Camoco identifies genes with known roles in elemental accumulation

To explore the broader biological processes represented among HPO genes, we performed GO enrichment analysis on the candidate lists, revealing enrichments for five elements (Supp. Table 10). For example, Sr was enriched for genes involved in anion transport (GO:0006820; *p* ≤ 0.008) and metal ion transmembrane transporter activity (GO:0046873; *p* ≤ 0.015) (See Supplementary Text for in-depth summary). Possibly due to insufficient functional annotation of the maize genome, these enrichment results were limited, and zero elements passed a strict multiple-test correction (Bonferroni). We created a larger set of genes including genes highly connected to the HPO genes, and compared those to GO terms (Supplementary Text). As detailed in the supplemental materials, several GO terms were enriched in this set and include genes that act in previously described pathways known to impact elemental traits (Supp. Figure 8; Supp. Table 11). However, GO terms were too broad or insufficiently specific to distinguish causal genes.

We also manually examined literature support for the association of candidate genes with ionomic traits (see Supplementary Text for in-depth summary). Complementing genes with known roles in elemental homeostasis, HPO gene sets for some ionomic traits included multiple genes encoding known members of the same pathway or protein complex. For example, one gene with highly pleiotropic effects on the maize kernel ionome is *sugary1* (*su1*; GRMZM2G138060) (Baxter et al., 2014) which was present among the HPO genes for Se accumulation (Supp. Table 7) based on the root co-expression network (ZmRoot-Se) but was linked to significant NAM GWAS SNPs for the elements P, K, and As. Previous analysis of lines segregating *su1* allele demonstrated effects on the levels of P, S, K, Ca, Mn, Fe, As, Se, and Rb in the seed. A number of transporters with known roles in ionome homeostasis were also identified among the HPO genes. Among these were a P-type ATPase transporter of the ACA P2B subfamily 4 (GRMZM2G140328; ZmRoot-Sr) encoding a homolog of known plasma membrane localized Ca transporters in multiple species (Baxter, 2003), an ABC transporter homolog of the family involved in organic acid secretion in the roots from the As HPO set (GRMZM2G415529; ZmRoot-As) (Badri et al., 2007), and a pyrophosphate energized pump (GRMZM2G090718; ZmPAN-Cd). These candidates suggest that biological signal was enriched by combining co-expression with GWAS and provided evidence of associations between multiple pathways and elemental homeostasis.

### Mutant analysis validates GA-signaling DELLA domain transcription factors influence the maize ionome

One of the high-confidence candidate genes, which appeared in the HPO sets comparing Cd and the ZmRoot network, is the gibberellin (GA)-signaling component and DELLA and GRAS domain transcription factor *dwarf9* (GRMZM2G024973; *d9* (Winkler and Freeling, 1994)). *d9* is one of two DELLA paralogs in the maize genome, the other being *dwarf8* (GRMZM2G144744; *d8*); both can be mutated to dominant-negative forms that display dwarf phenotypes and dramatic suppression of GA responses (Lawit et al., 2010). Camoco ranked *d9* among the high-confidence candidates for Cd but not *d8*, though both are present in the root-based co-expression network (ZmRoot). In the ZmRoot network, D9 was strongly co-expressed with 38 other HPO genes (Figure 7; See Supplementary Text). There was only moderate, but positive, co-expression between D8 and D9 transcripts (ZmRoot: z = 1.03; ZmPAN: z = 1.04). Given the indistinguishable phenotypes of the known dominant mutants of *d8* and *d9*, the most likely explanation for this result is that there was allelic variation for *d9* but not *d8* in the GWAS panel. Given the indistinguishable phenotypes of the known dominant mutants of d8 and d9, the most likely explanation for this result is that there was allelic variation for d9 but not d8 in the GWAS panel. These results suggested that GA signaling in the roots might shape the ionome and alters the accumulation of Cd in seeds, with potential impacts on human health.

**Figure 7.**
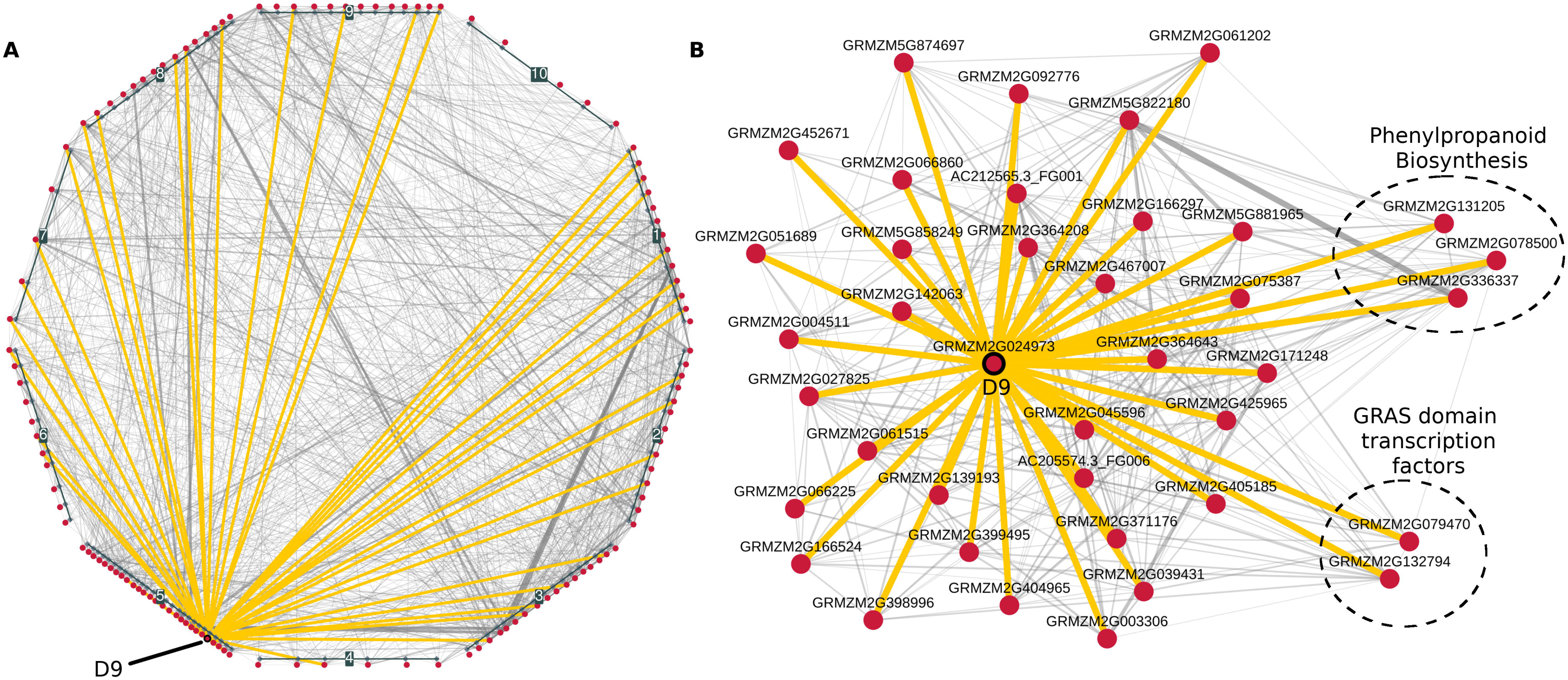
Co-expression network for D9 and cadmium HPO genes. Co-expression interactions among high-priority candidate (HPO) genes were identified in the ZmRoot network for Cd and visualized at several levels. Panel **(A)** shows local interactions among the 126 cadmium HPO genes (red nodes). Genes are grouped and positioned based on chromosomal location. Interactions among HPO genes and D9 (GRMZM2G024973) are highlighted in yellow. Panel **(B)** shows a force-directed layout of *D9* with HPO neighbors. Circled genes show sets of genes with previously known roles in elemental accumulation.

To test for an impact of GA signaling on the ionome and provide single-locus tests, we grew two dominant GA-insensitive mutants *D9-1* and *D8-mpl* and their congenic wild-type siblings (*sib9* and *sib8*). The dominant *D8-mpl* and *D9-1* alleles have nearly equivalent effects on above-ground plant growth and similar GA insensitivity phenotypes in the shoots (Winkler and Freeling, 1994). Both mutants were obtained from the maize genetics co-op and crossed three times to inbred B73 to generate BC2F1 families segregating 1:1 for the dwarf phenotype. Ears from phenotypically dwarf and phenotypically wild-type siblings were collected and processed for single-seed ionomic profiling using ICP-MS (Figure 8). Both dwarf lines had significantly different elemental compositions compared to their wild-type siblings. A joint analysis by *t*-tests between least-squared means comparing dwarfs and wild-types revealed that Cu, Fe, P, and Sr were higher in the dwarf than wild-type seeds (designated with two asterisks in Figure 8). Transcripts encoded by *d8* are expressed at lower levels than *d9* in the root but at many fold higher levels in the shoot (Wang et al., 2009; QTeller, 2018). *D8-mpl* was also significantly different from its sibling in Cd and Mo accumulation. It is possible that *D8-mpl* has a shoot-driven effect on Mo accumulation in the seed, but we note that previous work (Asaro et al., 2016) identified a large-effect QTL affecting Mo and containing the *mot1* gene a mere 22 Mb away from *d8*. As the allele at *mot1* is uncharacterized in the original *D8-mpl* genetic background, linkage drag carrying a *mot1* allele cannot be ruled out. The other dominant-negative allele, D9-1, did not recapitulate the Cd accumulation effect of the linked GWAS QTL that was the basis for its discovery as a high-confidence candidate gene by Camoco. However, the *D8-mpl* allele did recapitulate the accumulation effect, and our data demonstrate that both D8 and D9 have broad effects on other ionomic phenotypes.

**Figure 8.**
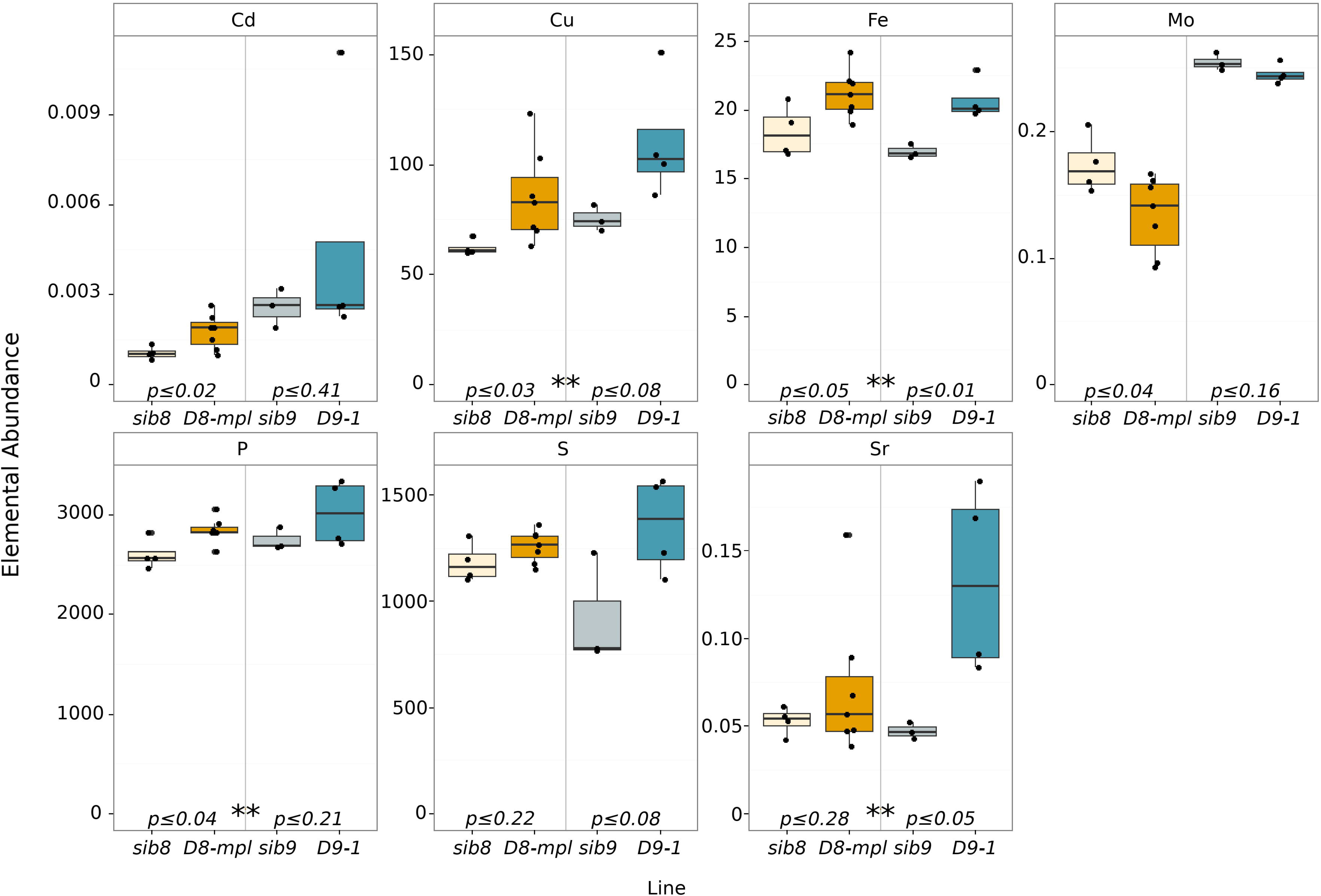
Ionomic profiles of *D8-mpl* and *D9-1* mutants. Box plots displaying ICP-MS values for *D8-mpl* and *D9-1* along with congenic wild-type siblings (*sib8* and *sib9*). Embedded p-values indicate statistical differences between mutants and wild-type siblings, while asterisks (**) indicate significant differences in a joint analysis between dwarf and wild-type.

Genes co-expressed with D9 that have annotated functions were investigated to determine which were further associated with ionomic traits, in particular, seed Cd levels (See Supplementary Text for in-depth report). Genes linked to the cell cycle, root development and Fe uptake suggest the hypothesis that maize DELLA-domain transcription factors regulate root architecture or the type II iron uptake mechanism used by grasses to affect the maize ionome.

### Camoco produce high-confidence candidate genes on large collection of non-ionomic GWAS

To assess the generalizability of our approach, we applied it to a separate collection of GWA studies surveying a compendium of phenotypes using the maize NAM population (Supp. Table 12). Using Camoco, SNPs were mapped to genes using two different window sizes (50kb and 100kb) and two flanking gene limits (1 and 2 genes). Gene-specific density and locality were calculated for each trait in all three co-expression networks, and HPO genes were identified as genes with less than 10% FDR in at least two SNP-to-gene mappings. Between 0 (Fructose, Leaf Length, Malate, Northern Leaf Blight, PCA of Metabolites PC2, Protein, Stalk Strength, and Total amino acid) and 302 (Average internode length (below ear)) HPO genes were discovered for the 41 traits examined (Supp. Table 12), with candidates produced for 33 of the 41 traits (80%). The candidate genes prioritized for these traits were largely non-overlapping with those discovered for the ionome traits: only 14 of 697 possible trait pairings (2%) overlapped significantly in terms of the candidate gene sets (Bonferroni corrected p-value < 0.05, Supp. Table 13). As with our maize ionome Camoco results, the genotype networks (ZmPAN and ZmRoot) outperformed the single accession map network (ZmSAM), supporting our earlier conclusion that genotypically diverse tissue networks support stronger candidate gene discovery for interpreting GWAS than tissue atlases. A full list of Wallace HPO genes can be found in Supp. Table 14.

## Discussion

Our approach addresses a challenging bottleneck in the process of translating large sets of statistically associated loci into shorter lists based on a more mechanistic understanding of these traits. Marker SNPs identified by a GWAS provide an initial lead on a region of interest, but due to linkage disequilibrium, the candidate region can be quite broad and implicate many potentially causal genes. In addition to LD, many SNPs identified by GWAS studies lie in regulatory regions quite far from their target genes (Clark et al., 2006; Castelletti et al., 2014;Louwers et al., 2009). Previous studies in maize found that while LD decays rapidly in maize (~1 kb), the variance can be large due to the functional allele segregating in a small number of lines (Wallace et al., 2014). Additionally, Wallace et al. showed that the causal polymorphism is likely to reside in regulatory regions, that is, outside of exonic regions (Wallace et al., 2014).

Relying solely on window based SNP-to-gene mapping can result in a very large (here, upward of 57% of all genes) and ambiguous set of candidate genes. Until we precisely understand the regulatory landscape in the species being studied, even the most powerful GWAS will identify polymorphisms that implicate genes many base pairs away. Here, we surveyed several different SNP-to-gene parameters, finding that the large majority of HPO genes were often not the closest genes to the identified SNPs (Supp. Figure 9). These genes would likely not have been identified using the common approach of prioritizing the genes closest to each marker SNP.

A common approach to interpreting lists of significant SNPs is through manual inspection of the genome region of interest with a goal of identifying candidate genes whose function is consistent with the phenotype of interest. This can introduce bias into the discovery process and necessarily ignores uncharacterized genes. For non-human and non-model species, like maize, this manual approach is especially ineffective because the large majority of the genome remains functionally under-characterized. Functional validation is expensive and time consuming. Combining data-driven approaches such as network integration with expert biological curation is an efficient means for the prioritization of genes driving complex traits like elemental accumulation, so that functional validation can be applied to only those best candidates. Camoco leverages orthogonal gene expression data, which can now be readily collected for most species of interest, to add an additional layer of relevant biological context to the interpretation of GWAS data and the prioritization of potentially causal variants for further experimental validation. In this way, Camoco complements approaches taken in model organisms and humans where probabilistic functional gene networks have been used to analyze GWAS datasets (Lee et al., 2010; Shim et al., 2017; Lee and Lee, 2018). Using RNA-Seq or other high throughput sequencing methods, high quality functional networks can be readily used in species without Bayesian networks. We evaluated our framework under simulated conditions as well as applied to a large scale GWAS in order to define different co-expression metrics and networks, biases such as *cis* co-expression, and network parameters needed to be considered in order to identify co-expression signal.

Camoco successfully identified subsets of genes linked to candidate SNPs that also exhibit strong co-expression with genes near other candidate SNPs. Integrating GWAS data with co-expression networks resulted a set of 610 HPO genes that are primed for functional validation (1.5% of the maize FGS). The resulting prioritized gene sets reflect groups of co-regulated genes that can potentially be used to infer a broader biological process in which genetic variation affects the phenotype of interest. Indeed, using Camoco, we found strong evidence for HPO gene sets in 13 of the 17 elemental accumulation phenotypes we examined (with 5 or more HPO genes). These high-priority sets of genes represent a small, high-confidence subset of the candidates implicated by the GWAS for each phenotype (see Supp. Table 6 and Figure 6).

It is important to note caveats of our approach. For example, phenotypes caused by genetic variation in a single or small number of genes or, alternatively, caused by a diverse set of otherwise functionally unrelated genes are not good candidates for our approach. The core assumption underpinning Camoco is that there are multiple variants in different genes, each driving phenotypic variation by virtue of their involvement in a common biological process. We expect that this assumption holds for many phenotypes (supported by the fact that we have discovered strong candidates for the most traits examined here), but we expect there are exceptional traits and causal genes that will violate this assumption. For these traits and genes, Camoco will not perform well.

Co-expression among genes can also arise due to processes unrelated to the GWAS trait being examined. For example, uncontrolled population structure in the samples used to construct the network can cause co-expression among genes which will introduce noise to overlap metrics. Camoco does correct for some of these cases, such as co-inheritance of *cis* regulatory elements. However, population structure driving co-expression of physically unlinked genes near GWAS SNPs, will lead to false positive co-expression interactions.

Finally, expression data used to build networks do not fully overlap with genomic data included in GWAS. For example, of the 39,656 genes in the maize filtered gene set, 11,718 genes did not pass quality control filters and were absent from the three co-expression networks analyzed here; they thus could not be analyzed despite the possibility there were potentially significant GWAS SNPs nearby.

### Relationship between Camoco and previous tools for GWAS analysis

It is important to note that previous studies have leveraged the complementarity of gene expression and/or other functional genomic data to interpret GWAS. For example, one powerful previously described approach is GWAB (Lee and Lee, 2018; Shim et al., 2017; Lee et al., 2011), which integrates functional networks and GWAS results to prioritize candidate genes, with applications described in Arabidopsis and human. These manuscripts focus on the use of integrated functional networks, which incorporate data from a diverse set of sources (e.g. protein-protein interaction networks, phylogenetic similarity, sequence similarity). Such networks have been built for Arabidopsis and human (and several other “data-rich” species), but their construction is not possible in many plant species where functional genomic data beyond expression simply does not exist. Here, we focus exclusively on co-expression networks as the basis for GWAS interpretation as these can be built for the majority of species where research communities are performing GWAS (because gene expression compendia have already been produced, or can be readily produced).

Another series of papers describe the use of co-expression networks from ATTED-II to interpret GWAS results in Arabidopsis (Chan et al., 2011; Corwin et al., 2016). There are two notable distinctions between our work and these studies. First, these papers focus on analyzing SNPs very near or within coding regions of genes (< 1kb for Chan et al., 2 significant SNPs in coding region for Corwin et al.). Here, we provide evidence for many traits where the co-expression network clustering of causal candidate genes is much stronger when one considers genes encoded quite far (e.g. > 100kb) from the associated SNPs, including genes that are not directly adjacent. Second, both of these studies leverage a single co-expression network from the ATTED-II database. Here, we explore the important issue of which gene expression data provides the most informative context for GWAS candidate gene prioritization (tissue/developmental assays vs. profiling of diverse individuals).

We note that there has also been previous work integrating co-expression networks with GWA studies, focused on interpreting human traits (Baillie et al., 2018; Bunyavanich et al., 2014;Calabrese et al., 2017). Most of these studies first cluster the co-expression network using no GWAS information, define modules, and then assess overlap between GWAS-identified loci and these modules. These studies are generally less focused on prioritizing individual candidate causal genes, and instead focus on characterizing broad modules with connections to traits of interest.

Our study explores several important issues affecting the integration of co-expression and GWAS results, provides new insights about best practices, and importantly, we provide a complete, scalable computational pipeline for constructing co-expression networks and GWAS results integration, which can be used in many different species as long gene expression data are available.

### Camoco-discovered gene sets are as coherent as GO terms

In evaluating the expected performance of our approach, we simulated the effect of imperfect SNP-to-gene mapping by assuming that GO terms were identified by a simulated GWAS trait. Neighboring genes (encoded nearby on the genome) were added to simulate the scenario where we could not resolve the causal gene from linked neighboring genes. This analysis was useful as it established the boundaries of possibility for our approach, that is, how much noise in terms of false candidate genes can be tolerated before our approach fails. As described in Figure 5, this analysis suggests a sensitivity of ~40% using a ±500-kb window to map SNPs to genes (two flanking genes maximum), or a tolerance of nearly 75% false candidates due to SNP-to-gene mapping. Therefore, if linkage regions implicated by GWAS extend so far as to include more than 75% false candidates, we would not be likely to discover processes as coherent as GO terms.

At the same window/flank parameter setting noted above, we were able to make significant discoveries (genes with FDR ≤ 0.30) for 7 of 17 elements (41%) using the density metric in the ZmRoot network. This success rate is remarkably consistent with what was predicted by our GO simulations at the same window/flanking gene parameter setting. Intriguingly, HPO gene sets alone were not significantly enriched for GO term genes, indicating that while the HPO gene sets and GO terms exhibited strikingly similar patterns of gene expression, the gene sets they described do not significantly overlap. It was not until the HPO gene sets were supplemented with co-expression neighbors that gene sets exhibited GO term enrichment, though the resulting terms were not very specific. We speculate that this is due to discovery bias in the GO annotations that were used for our evaluation, which were largely curated from model species and assigned to maize through orthology. There are likely a large number of maize-specific processes and phenotypes that are not yet characterized, yet have strong co-expression evidence and can be given functional annotations through GWAS.

Our analysis shows that loci implicated by ionomic GWAS loci exhibit patterns of co-expression as strong as many of the maize genes co-annotated to GO terms. Additionally, gene sets identified by Camoco have strong literature support for being involved in elemental accumulation despite not exhibiting GO enrichment. Indeed, one of the key motivations of our approach was that crop genomes like maize have limited species-specific gene ontologies, and this result emphasizes the extent of this limitation. Where current functional annotations, such as GO, rely highly on orthology, future curation schemes could rely on species-specific data obtained from GWAS and co-expression.

Beyond highlighting the challenges of a genome lacking precise functional annotation, these results also suggest an interesting direction for future work. Despite maize genes’ limited ontological annotations, many GWAS have been enabled by powerful mapping populations (e.g., NAM (McMullen et al., 2009)). Our results suggest that these sets of loci, combined with a proper mapping to the causal genes they represent using co-expression, could serve as a powerful resource for gene function characterization. Furthermore, our simulations using FCR indicate that researchers could use more permissive genome-wide significance cutoffs from GWAS as the networks act as robust filters against false positive genes. Systematic efforts to curate the results from such GWAS using Camoco and similar tools, then providing public access in convenient forms, would be worthwhile. Maize is exceptional in this regard due to its excellent genomic tools and powerful mapping populations. There are several other crop species with rich population genetic resources but limited genome functional annotation that could also benefit from this approach

### Co-expression context matters

Using our approach, we evaluated 17 ionomic traits for overlap with three different co-expression networks. Two of the co-expression networks were generated from gene expression profiles collected across a diverse set of individuals (ZmRoot, ZmPAN) and performed substantially better than the ZmSAM network, which was based on a large collection of expression profiles across different tissues and developmental stages derived from a single reference line (B73). We emphasize that this result is not a reflection of the data quality or even the general utility of the co-expression network used to derive the tissue/developmental atlas. Evaluations of this network showed a similar level of enrichment for co-expression relationships among genes involved in the same biological processes (Table 1) and had very similar network structure (Table 2). Instead, our results indicate that the underlying processes driving genotypic variation associated with traits captured by GWAS are better captured by transcriptional variation observed across genetically diverse individuals. Indeed, despite networks having similar levels of GO term enrichment (Table 1), the actual GO terms that drove that enrichment are quite different (Supp. Table 1), which is consistent with our previous analysis demonstrating that the experimental context of co-expression networks strongly influences which biological processes it captures (Schaefer et al., 2014).

Between the two co-expression networks based on expression variation across genotypically diverse individuals, we also observed differences depending on which tissues were profiled. Our co-expression network derived from sampling of root tissue across a diverse set of individuals (ZmRoot) provided the best performance at the FDR we analyzed (Figure 6), producing a total of 335 (326 from density and 11 from locality, 2 in both) HPO candidate genes as compared to 228 (all from locality) HPO candidate genes produced by the ZmPAN network, which was derived from expression profiles of whole seedlings. This result affirms our original motivation for collecting tissue-specific gene expression profiles: we expected that processes occurring in the roots would be central to elemental accumulation phenotypes, which were measured in kernels. However, the difference between the performance of these two networks was modest and much less significant than the difference between the developmental/tissue atlas-derived network and the diverse genotype-derived networks. Furthermore, we expect neither the ZmRoot nor the ZmPAN network to fully describe elemental accumulation processes. While ions are initially acquired from the soil via the root system, we do not directly observe their accumulation in the seed. The datasets presented here could be further complemented by additional tissue-specific data, such as genotypically diverse seed, stalk, or leaf networks.

The performance of the ZmRoot versus the ZmPAN network was also quite different depending on which network metric we used. Specifically, HPO gene discovery in the ZmRoot network was driven by the density metric, while performance of the ZmPAN network relied on the locality metric (Figure 6). Locality and density were positively correlated, but only modestly, in both networks (Supp. Figure 6), implying that these two metrics are likely complementary. Indeed, this relationship was also observed for density and locality of GO terms. Table 1 shows that both metrics had similar overall performance, each capturing ~40% of GO terms in each network; however, only ~25% was captured by both metrics, indicating that there are certain biological processes where one metric is more appropriate than the other. In addition to the tissue source differing between the ZmRoot and ZmPAN networks, the number of experimental accessions drastically differed as well (503 accessions in ZmPAN and 48 in ZmRoot), and this influenced the performance of network metrics. We showed that locality was sensitive to the number of accessions used to calculate co-expression (Supp. Table 8), which could partially explain the bias between network metrics and the number of input accessions. This result also suggests that the 46 accessions in ZmRoot did not saturate this approach for co-expression signal and that expanding the ZmRoot dataset to include more accessions would result in greater power to detect overlap and the identification of more true positives using the locality metric. In future work, it would be worthwhile to further understand the relationship between the network data source and which subnetwork metrics perform the best.

In general, our results strongly suggest that co-expression networks derived from expression experiments profiling genetically diverse individuals, as opposed to deep expression atlases derived from a single reference genotype, will be more powerful for interpreting candidate genetic loci identified in a GWAS. Furthermore, our findings suggest that where it is possible to identify relevant tissues for a phenotype of interest, tissue-specific expression profiling across genetically diverse individuals is an effective strategy. Identifying the best co-expression context for a given GWAS is an important consideration for data generation efforts in future studies.

## Methods

### Availability of data and material

Full GWAS information for all traits studied here are publically available from Ziegler et al. (Ziegler et al., 2017). FPKM values from RNA-Seq data for the ZmSAM network was used from Stelpflug et al. (Stelpflug et al., 2015). FPKM values for the ZmPAN network is available from Hirsch et al. (Hirsch et al., 2014). Raw RNASeq data used to build the ZmRoot network are available in NCBI BioProject PRJNA304663. All computer source code used in this study is available from http://www.github.com/schae234/Camoco.

### Software implementation of Camoco

Camoco (Co-analysis of molecular components) is a python library that includes a suite of command line tools to inter-relate and co-analyze different layers of genomic data. Specifically, it integrates genes present near GWAS loci with functional information derived from gene co-expression networks. Camoco was developed to build and analyze co-expression networks from gene transcript expression data (i.e., RNA-Seq), but it can also be utilized on other expression data such as metabolite, protein abundance, or microarray data.

This software implements three main routines: (1) construction and validation of co-expression networks from a counts or abundance matrix, (2) mapping SNPs (or other loci) to genes, and (3) an algorithm that assesses the *overlap* of co-expression among candidate genes near significant GWAS peaks.

Camoco is open source and freely available under the terms of the MIT license. Full source code, software examples, as well as instructions on how to install and run Camoco are available on GitHub (Camoco Software Repository, 2018). Camoco version 0.5.0 (DOI:10.5281/zenodo.1049133) was used for this article.

### Construction quality control of co-expression networks

#### Camoco Parameters

All networks were built (using the CLI) with the following Camoco QC parameters:

- min_expr_level: 0.001 (expression [FPKM] below this is set to NaN)
- max_gene_missing_data: 0.3 (genes missing expression data more than this percent were removed from analysis)
- max_accession_missing data: 0.08 (Accessions missing expression data in more than this percent were removed from analysis)
- min_single_sample_expr: 1.0 (genes must have at least this amount of expression [FPKM] in one accession)

#### ZmPAN: A genotypically diverse, PAN genome co-expression network

Camoco was used to process the fragments per kilobase per million reads (FPKM) table reported by Hirsh et al. and to build a co-expression network. The raw gene expression data were passed through the quality control pipeline in Camoco. After QC, 24,756 genes were used to build the network. For each pairwise combination of genes, a Pearson correlation coefficient (PCC) was calculated across FPKM profiles to produce ~306 million network edge scores (Supp. Figure 1A), which were then Fisher transformed and standard normalized (z-score hereafter) to allow cross network comparison (Supp. Figure 1B) (Huttenhower et al., 2006; Schaefer et al., 2014). A global significance threshold of z ≥ 3 was set on co-expression interactions in order to calculate gene degree and other conventional network measures.

To assess overall network health, several approaches were taken. First, the z-scores of edges between genes co-annotated in the maize gene ontology (GO) terms were compared to edges in 1,000 random terms containing the same number genes. Supp. Figure 1C shows the distribution of *p*-values compared to empirical z-score of edges within a GO term. With a nominal *p*-value cutoff of 0.05, the PAN co-expression network had 11.9-fold more GO terms than expected with *p* ≤ 0.05, suggesting that edges within this co-expression network capture meaningful biological variation. Degree distribution is also as expected within the network. Supp. Figure 1D shows empirical degree distributions compared to the power law, exponential, and truncated power law distributions. Typically, the degree distributions of biological networks are best fit by a truncated power law distribution, which is consistent with the ZmPAN genome co-expression network (Ghazalpour et al., 2006).

#### ZmSAM: A maize single accession map co-expression network

Publicly available gene expression data were generated from Stelpflug et al (Stelpflug et al., 2015). In total, 22,691 genes passed quality control metrics. Similar to the ZmPAN network described above, gene interactions were calculated between each pairwise combination of genes to produce ~257 million network edges. A global significance threshold of z ≥ 3 was set on co-expression interactions in order to differentiate significantly co-expressed gene pairs.

Supp. Figure 2A shows the distribution of edge scores before they were Fisher transformed and standard normalized (Supp. Figure 2B). The ZmSAM network shows a 10.8-fold enrichment for strong edge scores (*p* ≤ 0.05) between genes annotated to the same GO terms (Supp. Figure 2C). A final network health check shows that the empirical degree distribution of the ZmSAM network is consistent with previously characterized biological networks (Supp. Figure 2D).

#### ZmRoot: A genotypically diverse maize root co-expression network

Plants were grown from 48 diverse maize accessions: (A5554, B57, B73, B76, B97, CML103, CML108, CML157Q, CML158Q, CML228, CML277, CML311, CML322, CML341, CML69, CMl333, F2834T, F70NY2011, H84, H95 HP301, HY, IL14H, KY21, KY228, Ki11, Ki3, Ki44, M162W, M37W, MO17, MO18W, MS71, NC260, NC350, NC358, NC360, OH40B, OH43, OH7B, P39, SC357, T2I16, TX303, TZi8, U267Y, W22, W64A). Lines were selected to span a diverse panel starting with the 25 NAM parents, then adding more diverse lines that were at the extreme of accumulation for at least one element. 2-3 plants per genotype were distributed to independent trays and grown in the greenhouse soil mixture for 2 weeks and a 1-2 inch section of the root ~1 inch below the soil surface was collected and frozen in liquid nitrogen. Roots were ground in liquid nitrogen and RNA was extracted using Trizol. Sample quality was checked on a Bioanalyzer, and then 2 samples per genotype were pooled before library construction. Library construction and sequencing were done at the UMN sequencing core. RNA was extracted and sequenced in triplicate and multiplexed across 11 barcoded, multiplexed sequencing lanes using TruSeq Stranded RNA Library Prep and Illumina HiSeq 100-bp paired-end RNA sequencing (RNA-Seq) reads. Each library was split across two different Illumina HiSeq2000 lanes (between 6-10 lines multiplexed per lane) totaling 10 lanes with a final lane including all the libraries to help eliminate technical artifacts. Raw reads were deposited into the short read archive (SRA) under project number PRJNA304663.

Raw reads were passed through quality control using the program AdapterRemoval (Lindgreen, 2012), which collapses overlapping reads into high-quality single reads while also trimming residual PCR adapters. Reads were then mapped to the maize 5b reference genome using BWA (Li and Durbin, 2009; Schubert et al., 2014), PCR duplicates were detected and removed, and then realignment was performed across detected insertions and deletions, resulting in between 14 and 30 million high-quality, unique nuclear reads per accession. Two accessions (H84 and H95) were dropped due to low coverage, bringing the total number to 46.

Quantification of gene expression levels into FPKM was done using a modified version of HTSeq that quantifies both paired- and unpaired-end reads (Anders et al., 2014), available on GitHub (MixedHTSeq Software Repository, 2018). Raw FPKM tables were imported into Camoco and passed through the quality control pipeline. After QC steps, 25,260 genes were included in co-expression network construction containing ~319 million interactions. Supp. Figure 3A shows raw PCC scores, while Supp. Figure 3B shows z-scores after standard normal transformation. Similar to ZmPAN and ZmSAM, co-expression among GO terms was compared to random gene sets of the same size as GO terms (1,000 instances) showing a 13.5-fold enrichment for GO terms with significantly co-expressed genes (Supp. Figure 3C). The degree distribution of the ZmRoot network closely follows a truncated power law similar to the other networks built here (Supp. Figure 3D).

### SNP-to-gene mapping and effective loci

Two parameters are used during SNP-to-gene mapping: candidate window size and maximum number of flanking genes. Windows were calculated both upstream and downstream of input SNPs. SNPs having overlapping windows were collapsed down into *effective loci* containing the contiguous genomic intervals of all overlapping SNPs, including windows both upstream and downstream of the effective locus’ flanking SNPs (e.g., locus 2 in Figure 1A). Effective loci were cross referenced with the maize 5b filtered gene set (FGS) genome feature format (GFF) file (http://ftp.maizesequence.org/release-5b/filtered-set/ZmB73_5b_FGS.gff.gz) to convert effective loci to candidate gene sets containing all candidate genes within the interval of the effective SNP and also including up to a certain number of flanking genes both upstream and downstream from the effective SNP. For each candidate gene identified by an effective locus, the number of intervening genes was calculated from the middle of the candidate gene to the middle of the effective locus. Candidate genes were ranked by the absolute value of their distance to the center of their parental effective locus. Algorithms implementing SNP-to-gene mapping used here are accessible through the Camoco command line interface.

### Calculating subnetwork density and locality

Co-expression was measured among candidate genes using two metrics: density and locality. Subnetwork *density* is formulated as the average interaction strength between *all* (un-thresholded) pairwise combinations of input genes, normalized for the total number of input gene pairs:

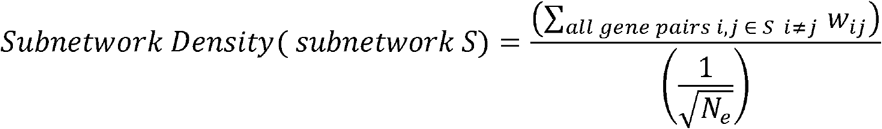

where *w_ij_* is the co-expression score between genes *i* and *j N_e_* is the number of total number of pairwise, non-self gene interactions in the subnetwork.

Network *locality* assesses the proportion of significant co-expression interactions (z ≥ 3) that are locally connected to other subnetwork genes compared to the number of global network interactions. To quantify network locality, both local and global degree are calculated for each gene within a subnetwork where local degree is the number of interactions to other genes in the subnetwork and global degree is the total number of interactions a gene has. To account for degree bias, where genes with a high global degree are more likely to have more local interactions, a linear regression is calculated on local degree using global degree (designated: local ~ global), and regression residuals for each gene are analyzed:

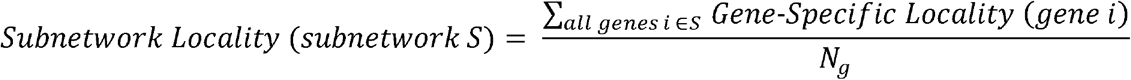

where the gene-specific locality measure is defined below (Eq. 4), and *N_g_* is the number of genes in the subnetwork of interest.

Gene-specific density is calculated by considering subnetwork interactions on a per-gene basis:

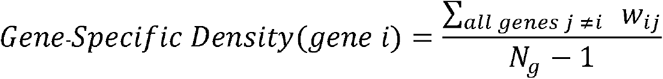

where *w_ij_* is the co-expression score between genes *i* and *j N_g_* is the total number of genes in the co-expression network.

Gene locality residuals can be interpreted independently to identify gene-specific locality:

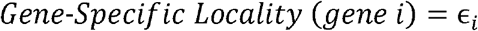

where ∊*_i_* is the residual for gene *i* derived from fitting the following regression model on the entire genome:

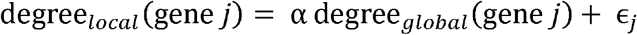

where degree_*local*_ (gene *j*) is the total number of interactions between gene *j* and the subnetwork of interest meeting the threshold, and degree_*global*_ (gene *j*) is the total number of interactions between gene *j* and any other gene in the genome.

Interactions among genes that originate from the same effective GWAS locus (i.e., *cis* interactions) were removed from density and locality calculations due to biases in *cis* co-expression. During SNP-to-gene mapping, candidate genes retained information containing a reference back to the parental GWAS SNP. A software flag within Camoco allows for interactions derived from the same parental SNP to be discarded from co-expression score calculations.

Statistical significance of subnetwork density and locality metrics (for both individual genes and whole subnetworks) was assessed by comparing the observed statistic to the distribution of 1,000 randomly sampled sets of candidate genes, conserving the number of input genes. This sampling was used to derive a null distribution, which was used to calculate an empirical p-value.

### Simulating GWAS using Gene Ontology (GO) terms

GO (Harris et al., 2004) annotations were downloaded for maize genes from http://ftp.maizesequenee.org/release-4a.53/funetional_annotations/. Co-annotated genes within a GO term were treated as true causal genes identified by a hypothetical GWAS. Terms between 50 and 100 genes were included to simulate the genetic architecture of a multi-genic trait. In each co-expression network, terms having genes with significant co-expression (*p*-value ≤ 0.05; density or locality) were retained for further analysis. Noise introduced by imperfect GWAS was simulated using two different methods to decompose how noise affects significantly co-expressed networks.

#### Missing Candidate Rate

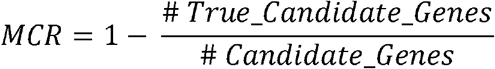

#### False Candidate Rate

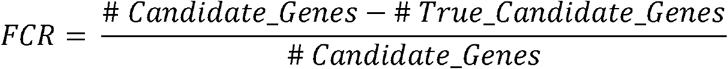

#### Simulating missing candidate gene rate (MCR)

The effects of MCR were evaluated by subjecting GO terms with significant co-expression (*p* ≤ 0.05; described above) to varying levels of missing candidate rates. True GO term genes were replaced with random genes at varying rates (MCR: 0%, 10%, 20%, 50%, 80%, 90%, 100%). The effect of MCR was evaluated by assessing the number of GO terms that retained significant co-expression (compared to 1,000 randomizations) at each level of MCR.

#### Adding false candidate genes by expanding SNP-to-gene mapping parameters

To determine how false candidates due to imperfect SNP-to-gene mapping affected the ability to detect co-expressed candidate genes linked to a GWAS trait, GO terms with significantly co-expressed genes were reassessed after incorporating false candidate genes. Each gene in a GO term was treated as a SNP and remapped to a set of candidate genes using the different SNP-to-gene mapping parameters (all combinations of 50 kb 100 kb, 500 kb and one, two, or five flanking genes). Effective FCR at each SNP-to-gene mapping parameter setting was calculated by dividing the number of true GO genes with candidates identified after SNP-to-gene mapping. Since varying SNP-to-gene mapping parameters changes the number of candidate genes considered within a term, each term was considered independently for each parameter combination.

### Maize ionome GWAS

Elemental concentrations were measured for 17 different elements in the maize kernel using inductively coupled plasma mass spectrometry (ICP-MS) as described in Ziegler et al. (Ziegler et al., 2017) Outliers were removed from single-seed measurements using median absolute deviation (Davies and Gather, 1993). Basic linear unbiased predictors (BLUPs) for each elemental concentration were calculated across different environments and used to estimate variance components (Hung et al., 2012). Joint-linkage analysis was run using TASSEL version 3.0 (Bradbury et al., 2007) with over 7,000 SNPs obtained by genotype by sequencing (GBS) (Elshire et al., 2011). An empirical *p*-value cutoff was determined by performing 1,000 permutations in which the BLUP phenotype data were shuffled within each NAM family before joint-linkage analysis was performed. The *p*-value corresponding to a 5% false discovery rate was used for inclusion of a QTL in the joint-linkage model.

Genome-wide association was performed using stepwise forward regression implemented in TASSEL version 4.0 similar to other studies (Wallace et al., 2014; Cook et al., 2012;Tian et al., 2011). Briefly, genome-wide association was performed on a chromosomal-by-chromosome basis. To account for variance explained by QTLs on other chromosomes, the phenotypes used were the residuals from each chromosome calculated from the joint-linkage model fit with all significant joint-linkage QTLs except those on the given chromosome. Association analysis for each trait was performed 100 times by randomly sampling, without replacement, 80% of the lines from each population.

The final input SNP dataset contained 28.9 million SNPs obtained from the maize HapMap1 (Gore et al., 2009), the maize HapMap2 (Chia et al., 2012), as well as an additional ~800,000 putative copy-number variants from analysis of read depth counts in HapMap2 (Wallace et al., 2014; Chia et al., 2012). These ~30 million markers were projected onto all 5,000 lines in the NAM population using low-density markers obtained through GBS. A cutoff *p*-value value (*p* ≤ 1e-6) was used from inclusion in the final model. SNPs associated with elemental concentrations were considered significant if they were selected in more than 5 of the 100 models (resample model inclusion probability [RMIP]) (Valdar et al., 2009).

### Identifying ionome high-priority overlap (HPO) genes and HPO+ genes

Gene-specific density and locality were calculated for candidate genes identified from the 17 ionome GWAS traits as well as for 1,000 random sets of genes of the same size. Gene-specific metrics were converted to the standard normal scale (z-score) by subtracting the average gene-specific score from the randomized set and dividing by the average randomized standard deviation. A false discovery rate was established by incrementally evaluating the number of GWAS candidates discovered at a z-score threshold compared to the average number discovered in the random sets. For example, if ten GWAS genes had a gene-specific z-score of 3 and an average of 2.5 randomized genes (in the 1,000 random sets) had a score of 3 or above, the FDR would be 25%.

High-priority overlap (HPO) candidate genes for each element were identified by requiring candidate genes to have a co-expression FDR ≤ 30% in two or more SNP-to-gene mapping scenarios in the same co-expression network using the same co-expression metric (i.e., density or locality).

HPO+ candidate gene sets were identified by taking the number of HPO genes discovered in each element (*n* genes) and querying each co-expression network for the set of (*n*) genes that had the strongest aggregate co-expression. For example, of the 18 HPO genes for P, an additional 18 genes (36 total) were added to the HPO+ set based on co-expression in each of the networks. Genes were added based on the sum of their co-expression to the original HPO set.

### Reduced-accession ZmPAN networks

Both the ZmPAN and ZmRoot networks were rebuilt using only the 20 accessions in common between the 503 ZmPAN and 46 ZmRoot experimental datasets. The ZmPAN network was also built using the common set of 20 accessions as well as 26 accessions selected from the broader set of 503 to simulate the number of accessions used in the ZmRoot network. Density and locality were assessed in these reduced-accession networks using the same approach as the full datasets.

### Identifying High Priority Genes from 41 non-Ionomic GWAS

Camoco was used to identify HPO candidate genes from 41 GWAS traits reported previously by Wallace et al. (Wallace et al., 2014) which included: 100 Kernel weight, Anthesis-silking interval, Average internode length (above ear), Average internode length (below ear), Average internode length (whole plant), Boxcox-transformed leaf angle, Chlorophyll A, Chlorophyll B, Cob diameter, Days to anthesis, Days to silk, Ear height, Ear row number, Fructose, Fumarate, Glucose, Glutamate, Height above ear, Height per day (until flowering), Leaf length, Leaf width, Malate, Nitrate, Nodes above ear, Nodes per plant, Nodes to ear, Northern Leaf Blight, PCA of metabolites: PC1, PCA of metabolites: PC2, Photoperiod Growing-degree days to silk, Photoperiod growing-degree days to anthesis, Plant height, Protein, Ratio of ear height to total height, Southern leaf blight, Stalk strength, Starch, Sucrose, Tassel branch number, Tassel length, Total amino acids. SNPs were mapped to genes using two window sizes (50kb and 100kb) as well as two flanking gene parameters (1 and 2 genes). Overlap was calculated using both density and locality in all three co-expression networks and FDR was calculated for candidate genes in each GWAS subnetwork as described above. High priority overlap (HPO) candidate genes were identified as described above as candidates genes with less than 10% FDR in at least two SNP-to-gene mappings (Supp. Table 12).

## Abbreviations

GWAS: Genome-wide Association Study
SNP: Single nucleotide polymorphism
LD: linkage disequilibrium
QTL: quantitative trait locus
Camoco: co-analysis of molecular components
FDR: false discovery rate
GO: Gene Ontology
MCL: Markov clustering algorithm
MCR: missing candidate rate
FCR: false candidate rate
ICP-MS: inductively coupled plasma mass spectrometry
NAM: nested association mapping
RMIP: resampling model inclusion probability
HPO: high priority overlap
ABC: ATP-binding cassette
RIL: recombinant inbred line
PRC: polycomb repressive complex
RBR: retinoblastoma-related proteins
CLI: command line interface
FPKM: fragments per kilobase per million reads
PCC: Pearson correlation coefficient
SRA: short read archive
FGS: filtered gene set
GFF: gene feature format
BLUP: Basic linear unbiased predictor
GBS: genotype by sequencing

## Acknowledgements

We would like to thank Ben VanderSluis, Henry Ward, and Joanna Dinsmore for their helpful comments and feedback in writing this article. We would also like to thank Abby Cabunoc-Mayes and other members of the Mozilla Science Lab for their mentorship and help in making Camoco a free and open scientific resource.

## Authors’ contributions

Experimental concept and design: CLM, OH, IB; Sample collection and data contribution: IB; Data analysis and interpretation: RS, JM, OH, BD, IB, CLM; Computational support: JJ; Manuscript writing and figures: RS, BD, IB, CLM; Manuscript review: All authors read and approved the final manuscript.

## Supplementary Tables

**Supp. Table 1**

**Full gene ontology term density and locality p-values.** Density and locality scores were measured between genes within each GO term. Subnetwork p-values were generated for both density and locality by comparing each term’s metric to 1,000 randomized gene sets of the same size. The “Number of Network Genes in GO Terms” indicates the intersect between genes present in the network and genes annotated to a GO term. Supports Table 1.

**Supp. Table 2**

**Network MCL cluster gene assignments.** Clusters in all three networks were identified using the MCL algorithm. Genes in each network were assigned to cluster IDs. Lower cluster IDs have a larger number of genes. Supports Table 2.

**Supp. Table 3**

**Network MCL cluster GO enrichment.** Enrichment of genes co-annotated for GO terms in each MCL cluster. Significance of enrichment was calculated using the hypergeometric test with a Bonferroni corrected p-value of ≤ 0.05. Supports Table 2.

**Supp. Table 4**

**Network signal of GO terms with various levels of MCR/FCR.** Co-expression among genes co-annotated to GO terms was compared to random gene sets of the same size to generate p-values. Noise was introduced by varying the missing candidate rate (MCR) or false candidate rate (FCR). Missing candidates were removed in proportion to the values in the table, while false candidates were introduced using SNP-to-gene mapping values (see WindowSize and FlankLimit columns). FCR values are reported as averages across 10% quantiles (see Figure 5). Supports Figure 4 and Figure 5.

**Supp. Table 5**

**Maize grain ionome SNP-to-gene mapping results.** Significant GWAS SNPs associated with the maize grain ionome were mapped to candidate genes. SNPs within overlapping windows were collapsed down to Effective Loci. Candidate genes were mapped by taking genes upstream and downstream (designated by Window Size) of the effective locus, up to the maximum designated by Flank Limit. The Ionome average shows the average per column for each value (e.g. at 50kb there are an average of 138 effective loci) as well as total average for the whole group (e.g. average of 50kb, 100kb, and 500kb is 119 effective loci). Supports Figure 6.

**Supp. Table 6**

**Maize grain ionome GWAS network overlap candidate genes.** Candidate genes were identified in each Co-expression network (ZmSAM, ZmPAN, or ZmRoot) using SNP-to-gene mapping for each element (using WindowSize and FlankLimit). Co-expression (density or locality) among all genes within a subnetwork was compared to randomized gene sets of the same size to establish p-values. Gene-specific z-scores were computed by comparing the empirical gene-specific density (Eq. 3) or locality (Eq. 4) to the average density or locality observed in randomized gene sets, then correcting for standard deviation. False discovery rates (FDRs) were calculated for candidate genes with positive gene-specific Co-expression values by comparing the number of genes discovered at a z-score cutoff to the average number of genes discovered in randomized sets. Supports Figure 6.

**Supp. Table 7**

**Maize grain ionome GWAS high-priority overlap (HPO) candidate genes.** High-priority overlap (HPO) genes were identified by calculating gene-specific density or locality (Method column) for each element at different SNP-to-gene mapping parameters (see WindowSize and FlankLimit columns). At an FDR cutoff of 30%, genes were defined as HPO if they were observed at two or more SNP-to-gene mapping parameters. Supports Figure 6.

**Supp. Table 8**

**Locality HPO genes discovered with networks built from accessions subsets.** The number of HPO genes discovered in full ZmPAN (503 accessions) and ZmRoot (46 accessions) networks was compared to networks built with a subset of accessions. Both ZmPAN and ZmRoot networks were re-built using a common set of 20 accessions. The ZmPAN network was re-built using 46 accessions consisting of the 20 common accessions and either 26 random or 26 CML biased accessions to simulate the number used in the full 46 accession ZmRoot network. Each network was analyzed for HPO genes in the 17 GWAS elements using locality. Supports Figure 6.

**Supp. Table 9**

**Multiple element HPO gene list.** The number of commonly discovered HPO genes, hypergeometric p-values of set overlap, and GRMZM IDs across multiple elements. Supports Figure 6.

**Supp. Table 10**

**Element gene ontology enrichment.** HPO genes for each element were tested for enrichment among genes co-annotated for gene ontology (GO) terms (hypergeometric test). Bonferroni correction is included as a column, treating each GO term as an independent test. Supports Figure 6.

**Supp. Table 11**

**HPO plus neighbors’ gene ontology enrichment.** Elemental HPO gene sets were supplemented with an additional set of highly connected neighbors equal to the number of genes in the HPO set. These HPO+ gene sets were tested for enrichment among genes annotated for GO terms (hypergeometric test). Supports Figure 6.

**Supp. Table 12**

**HPO genes discovered from non-ionomic traits.** HPO genes were identified with Camoco using SNPs from 41 GWAS described in Wallace et al. SNP-to-gene mapping was performed using 50kb and 100kb windows including either 1 or 2 additional flanking genes upstream and downstream of effective loci. Gene specific density and locality metrics for each trait were compared to (n=1,000) random sets of genes of the same size to establish a 10% FDR. Genes were considered HPO if they were observed in two or more SNP to gene mappings (see Materials and Methods). Candidates in the "Either" column are HPO genes discovered by either density or locality in any network. The number of genes discovered for each element is further broken down by Co-expression method (density, locality, both) and by network (ZmPAN, ZmSAM, ZmRoot). Candidates in the “Both” column were discovered by density and locality in the same network or in different networks (Any). Note: zero elements had HPO genes using “Both” methods in the ZmPAN and ZmSAM networks. Supports Figure 1.

**Supp. Table 13**

**Overlap between Wallace et al and ionome HPO genes.** Wallace and the ionome were compared for overlap between HPO using the hypergeometric distribution. P-values and accompanying Bonferroni indicate if the genes common between the GWAS traits are statistically significant. Supports Supp. Table 12.

**Supp. Table 14**

**ZmWallace GWAS network overlap candidate genes.** Candidate genes were identified in each Co-expression network (ZmPAN, ZmSAM, ZmRoot) using SNP-to-gene mapping for each GWAS (using Window Size and Flank Limit). Co-expression (density or locality) among all genes within a subnetwork was compared to randomized gene sets of the same size to establish subnetwork p-values. Gene specific z-scores were computed by comparing the empirical density (Eq. 3) or locality (Eq. 4) to the average density or locality observed in randomized gene sets, then correcting for standard deviation. False discovery rates (FDR) were calculated for candidate genes with positive gene-specific Co-expression values by comparing the number of genes discovered at a z-score cutoff to the average number of genes discovered in randomized sets. Supports Supp. Table 12.

## Supplementary Figures

**Supp. Figure 1.**
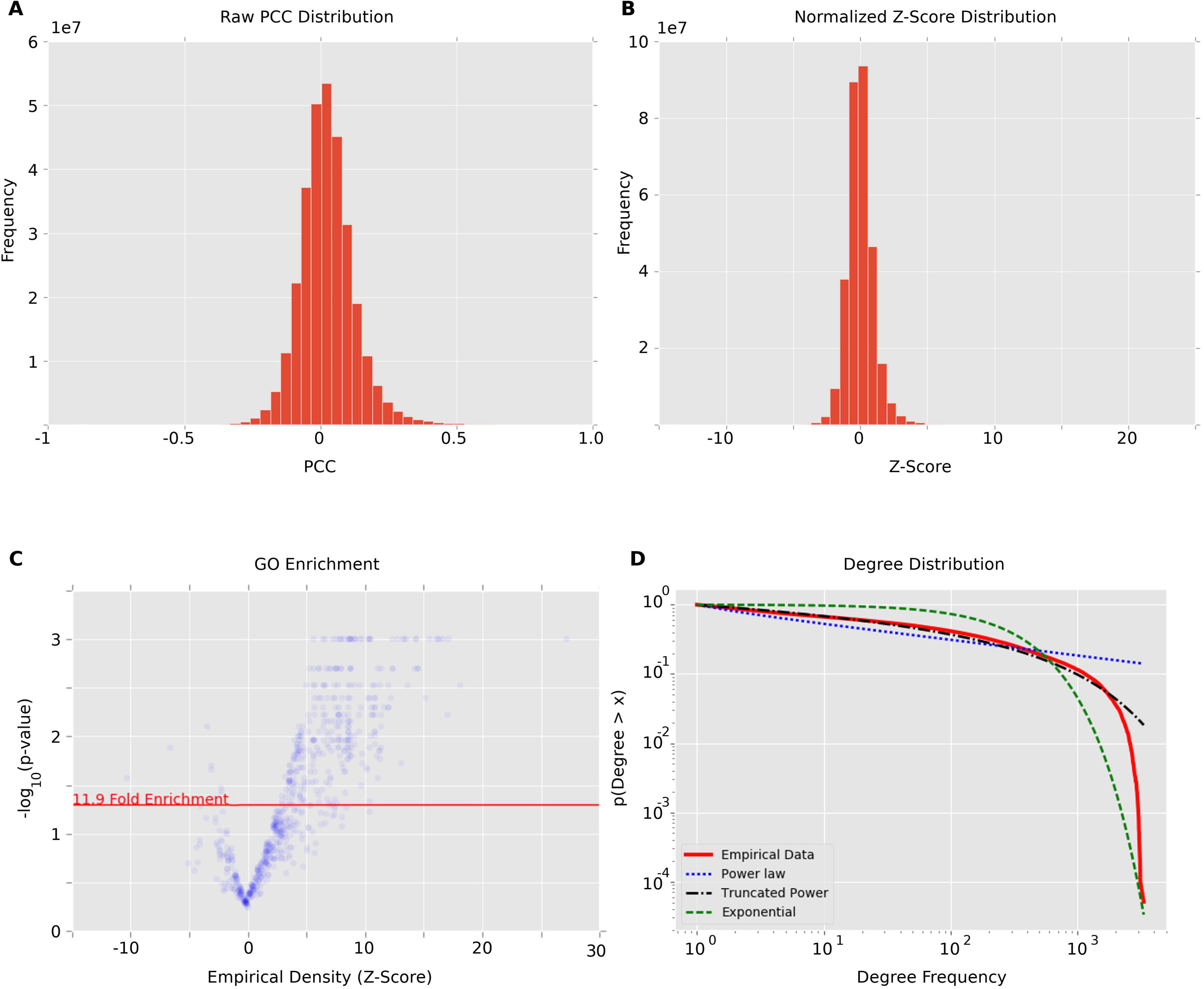
ZmPAN network health. Global network health of the maize PAN genome (ZmPAN) Co-expression network. **(A)** Raw Pearson correlation coefficient distribution of all Co-expression interactions. **(B)** Fisher-transformed, variance-stabilized, and mean centered network interactions. **(C)** A volcano plot showing empirical density for genes for each GO term compared to the corresponding *p*-value derived from measuring density in 1,000 random gene sets of the same size. Data points are transparent to show denseness. **(D)** Degree distribution of ZmPAN genome Co-expression network compared to power law, exponential, and truncated power law distributions. Supports Figure 1.

**Supp. Figure 2.**
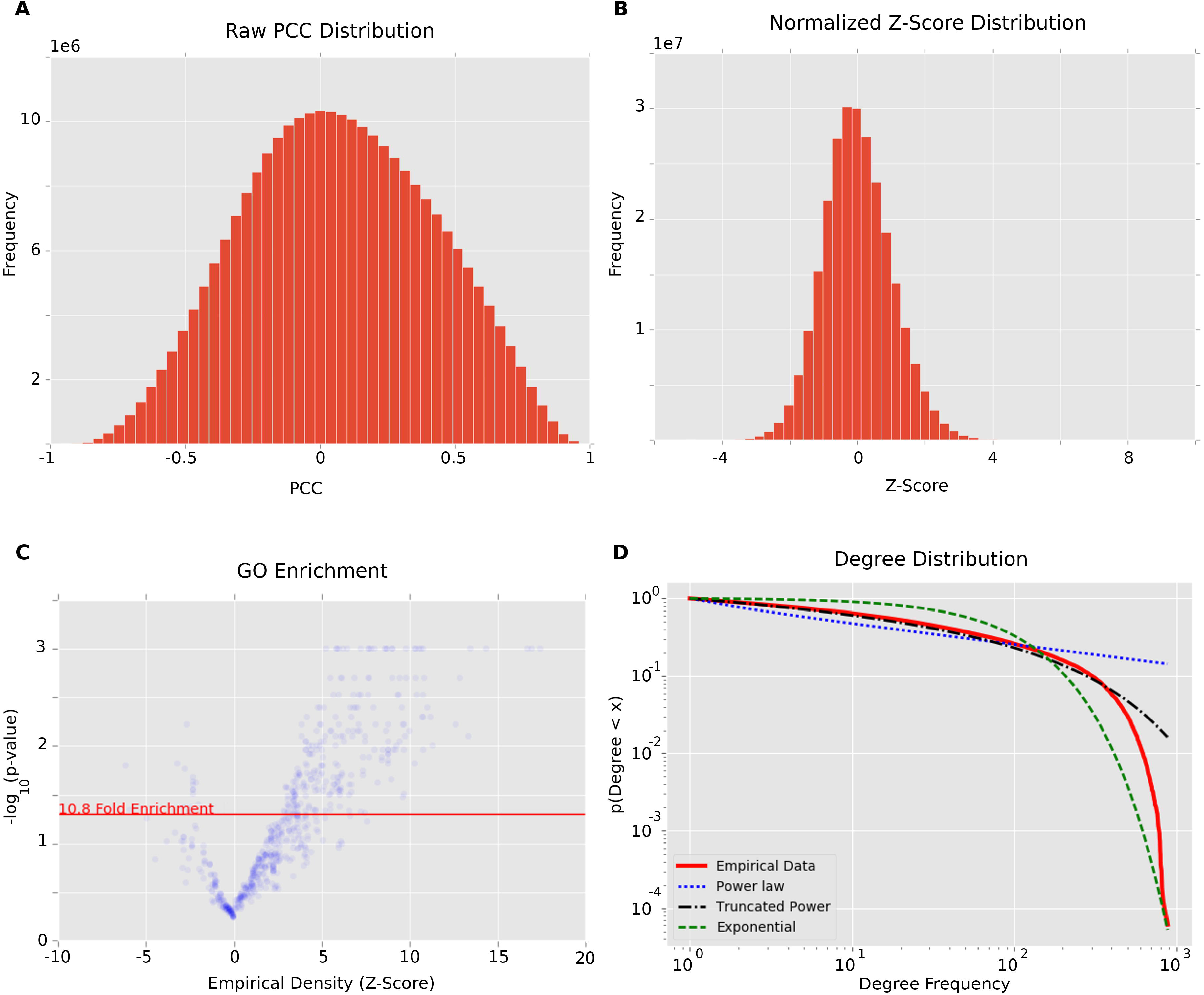
ZmSAM network health. Global network health of the maize ZmSAM Co-expression network. **(A)** Raw Pearson correlation coefficient distribution of all Co-expression interactions. **(B)** Variance-stabilized and mean centered network interactions. **(C)** A volcano plot showing empirical density for genes for each GO term compared to the corresponding *p*-value derived from measuring density in 1,000. random gene sets of the same size. Data points are transparent to show denseness. **(D)** Degree distribution of tissue/developmental Co-expression network compared to power law, exponential, and truncated power law distributions. Supports Figure 1.

**Supp. Figure 3.**
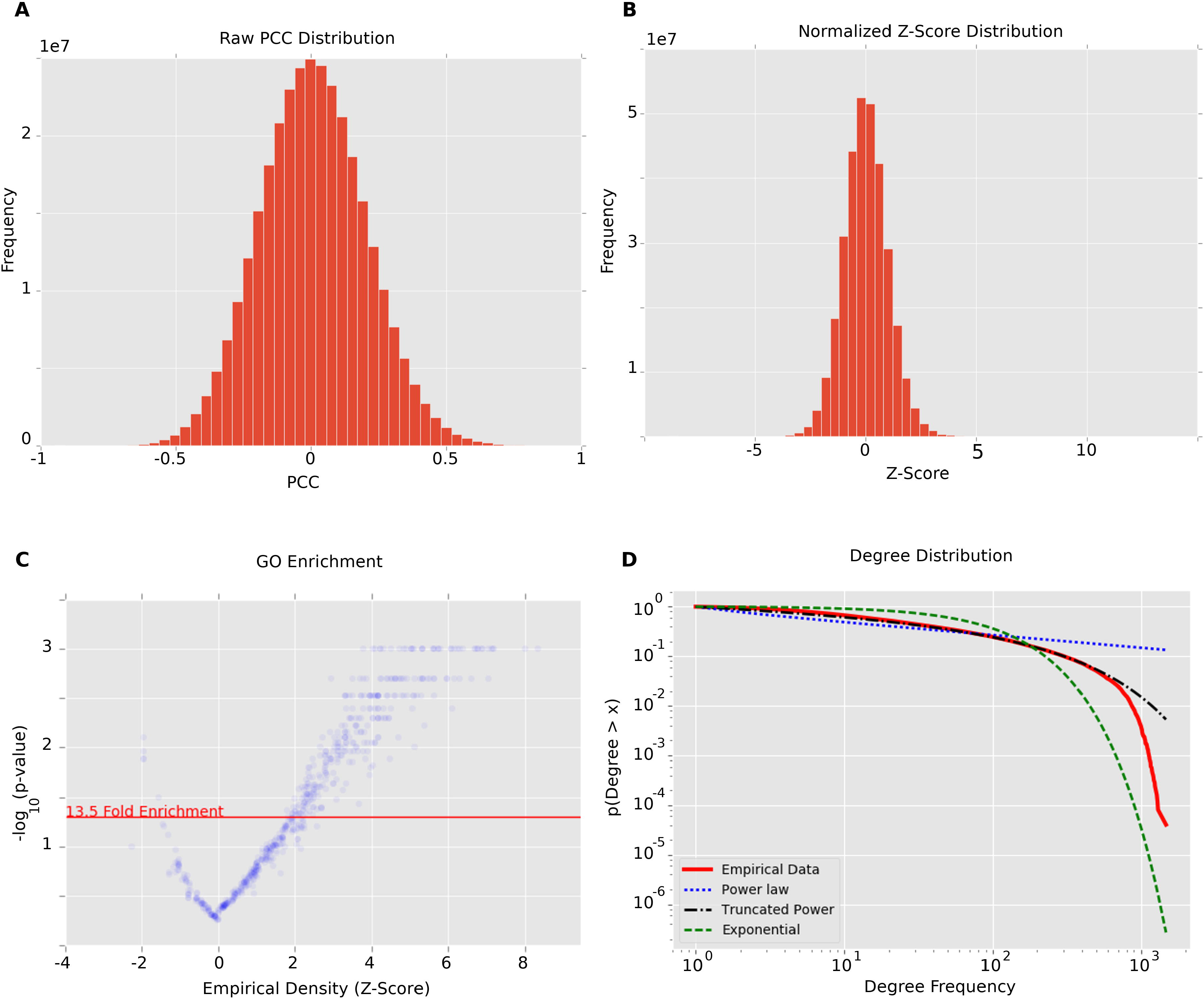
ZmRoot network health. Global network health of the maize ZmRoot Co-expression network. **(A)** Raw Pearson correlation coefficient distribution of all Co-expression interactions. **(B)** Variance-stabilized and mean centered network interactions. **(C)** A volcano plot showing empirical density for genes for each GO term compared to the corresponding *p*-value derived from measuring density in 1,000 random gene sets of the same size. Data points are transparent to show denseness. **(D)** Degree distribution of ZmRoot Co-expression network compared to power law, exponential, and truncated power law distributions. Supports Figure 1.

**Supp. Figure 4.**
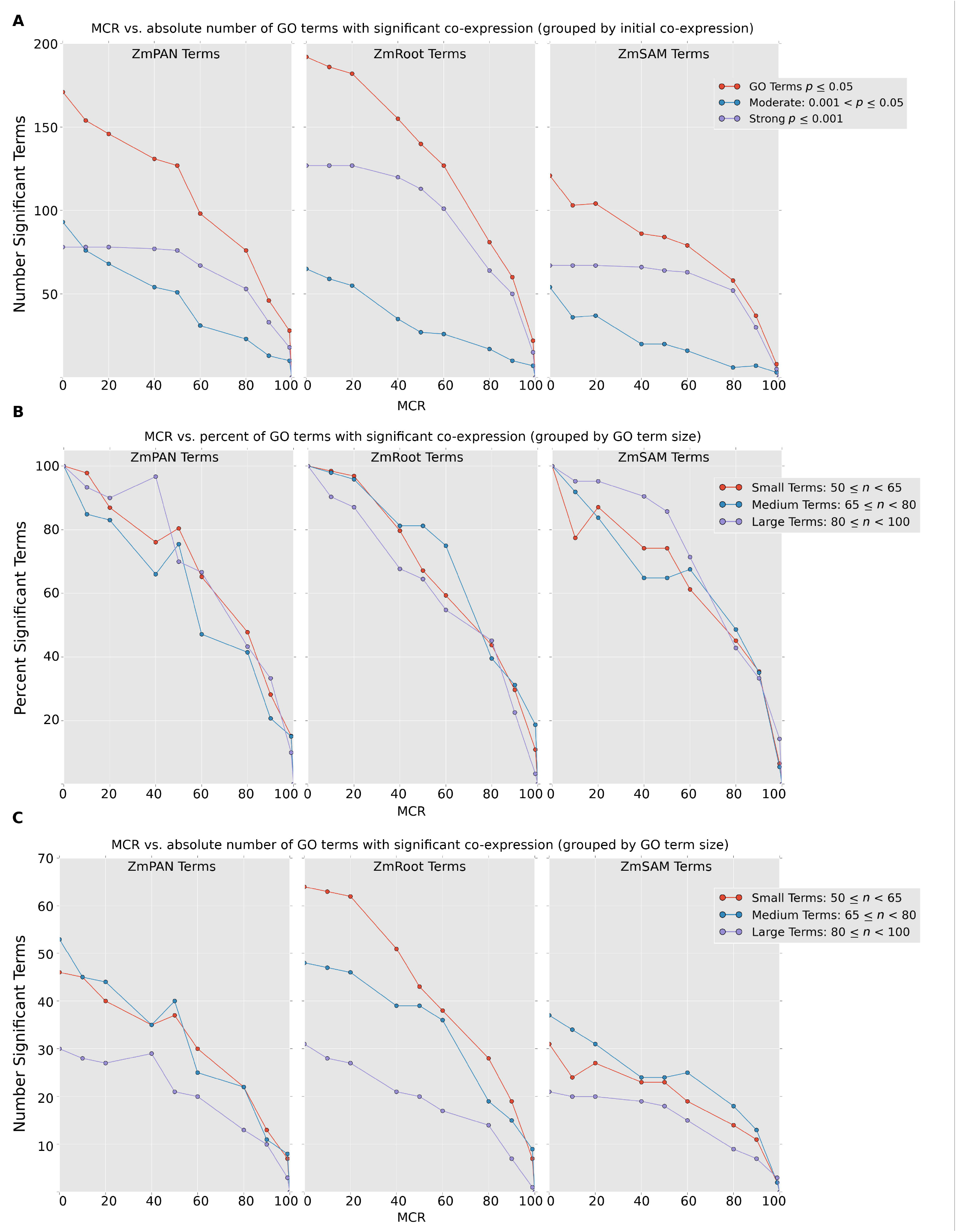
MCR supplemental figure. Panel **(A)** shows the absolute number of GO terms that remain significantly co-expressed at varying levels of MCR in each network. Red curves show all GO terms with an initial Co-expression *p*-value ≤ 0.05. Blue and violet curves show GO terms with either moderate or strong initial Co-expression (at MCR = 0). Panels **(B-C)** show the percent and absolute number of GO terms that remain significantly co-expressed at varying levels of MCR. The red curves show small GO terms (50 ≤ *n* < 65), the blue curves show medium sized GO terms (65 ≤ *n* < 80), and the violet curves show large terms (80 ≤ *n* < 100). Supports Figure 4.

**Supp. Figure 5.**
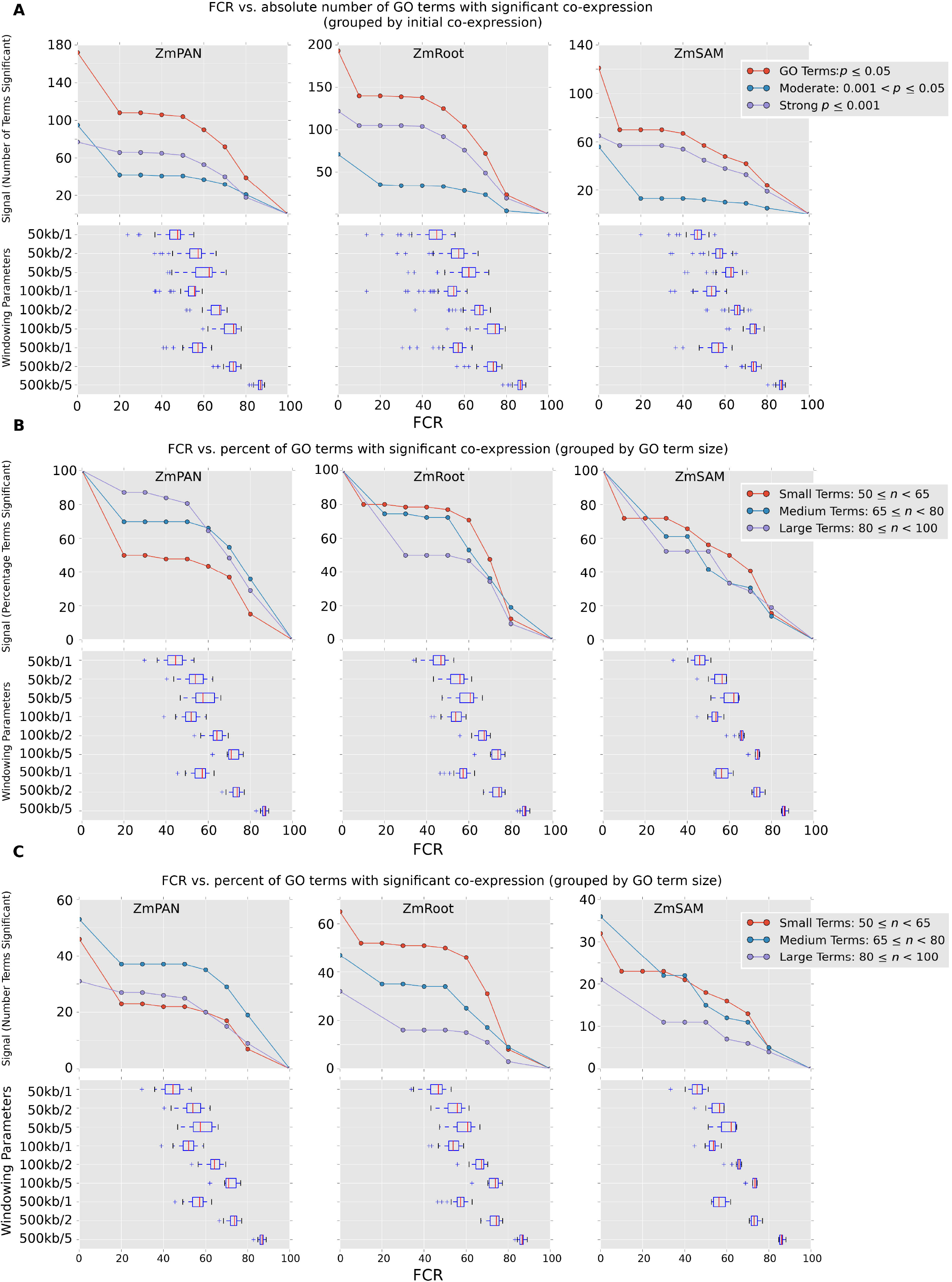
FCR supplemental figure. Panel **(A)** shows the absolute number of GO terms that remain significantly co-expressed at varying levels of FCR in each network. Red curves show all GO terms with an initial Co-expression *p*-value ≤ 0.05. Blue and violet curves show GO terms with either moderate or strong initial Co-expression. Panels **(B-C)** show the percent and absolute number of GO terms that remain significantly co-expressed at varying levels of FCR. The red curves show small GO terms (50 ≤ *n* < 65), the blue curves show medium sized GO terms (65 ≤ *n* < 80) and the violet curves show large terms (80 ≤ *n* < 100). Supports Figure 5.

**Supp. Figure 6.**
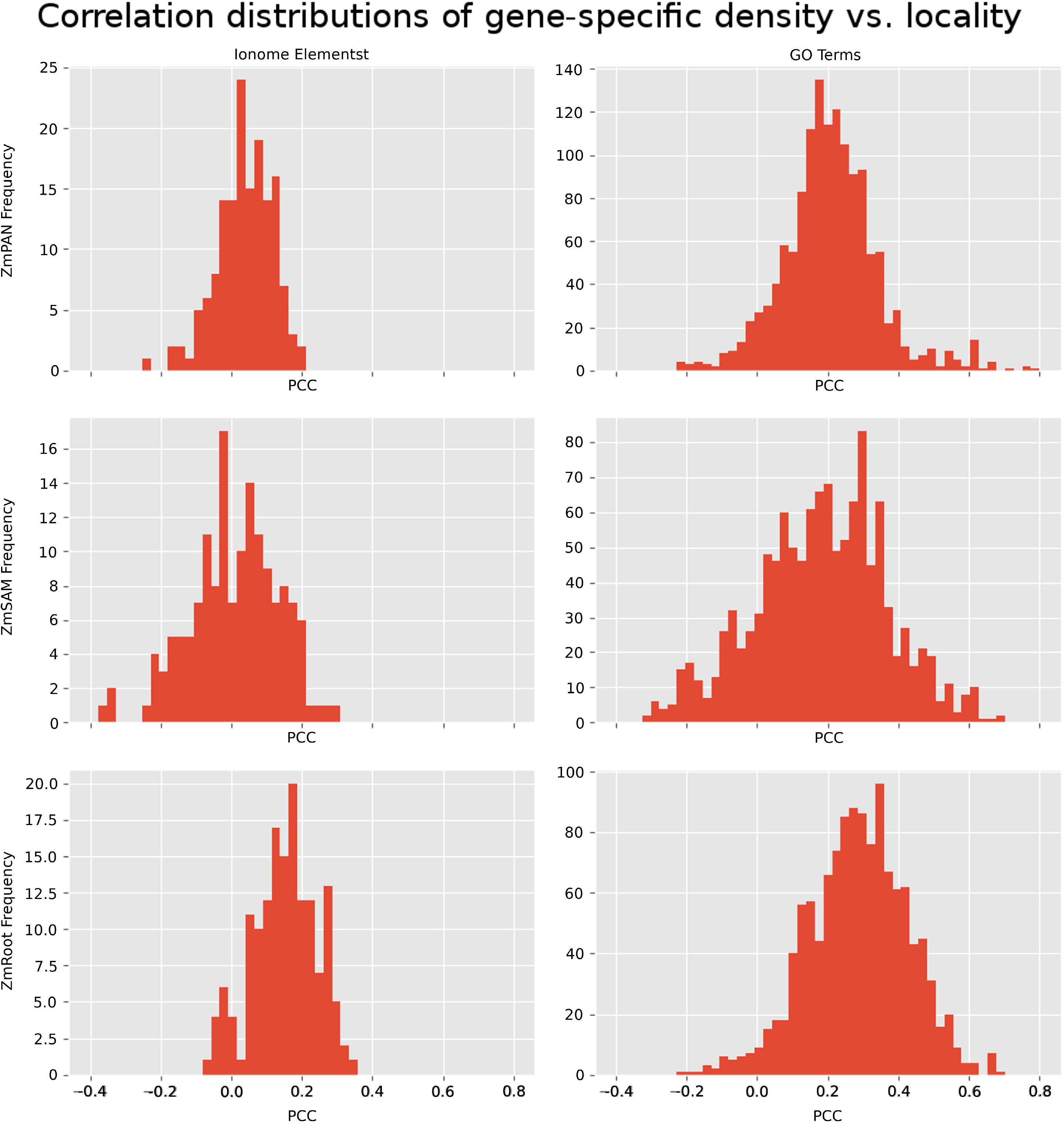
Distribution of Pearson correlation coefficients between gene-specific density and locality. Pearson correlation was measured between gene-specific density and locality in each network for both ionome elements and GO terms. PCCs between metrics were calculated by grouping sets of genes in either ionome elements (e.g., Al, Fe) or GO terms at the same SNP-to-gene mapping parameters (50-, 100-, and 500-kb window size and one, two, and five gene flank limits). The distribution shows the PCCs between the metrics aggregated across all SNP-to-gene mapping parameters. Supports Table 1.

**Supp. Figure 7.**
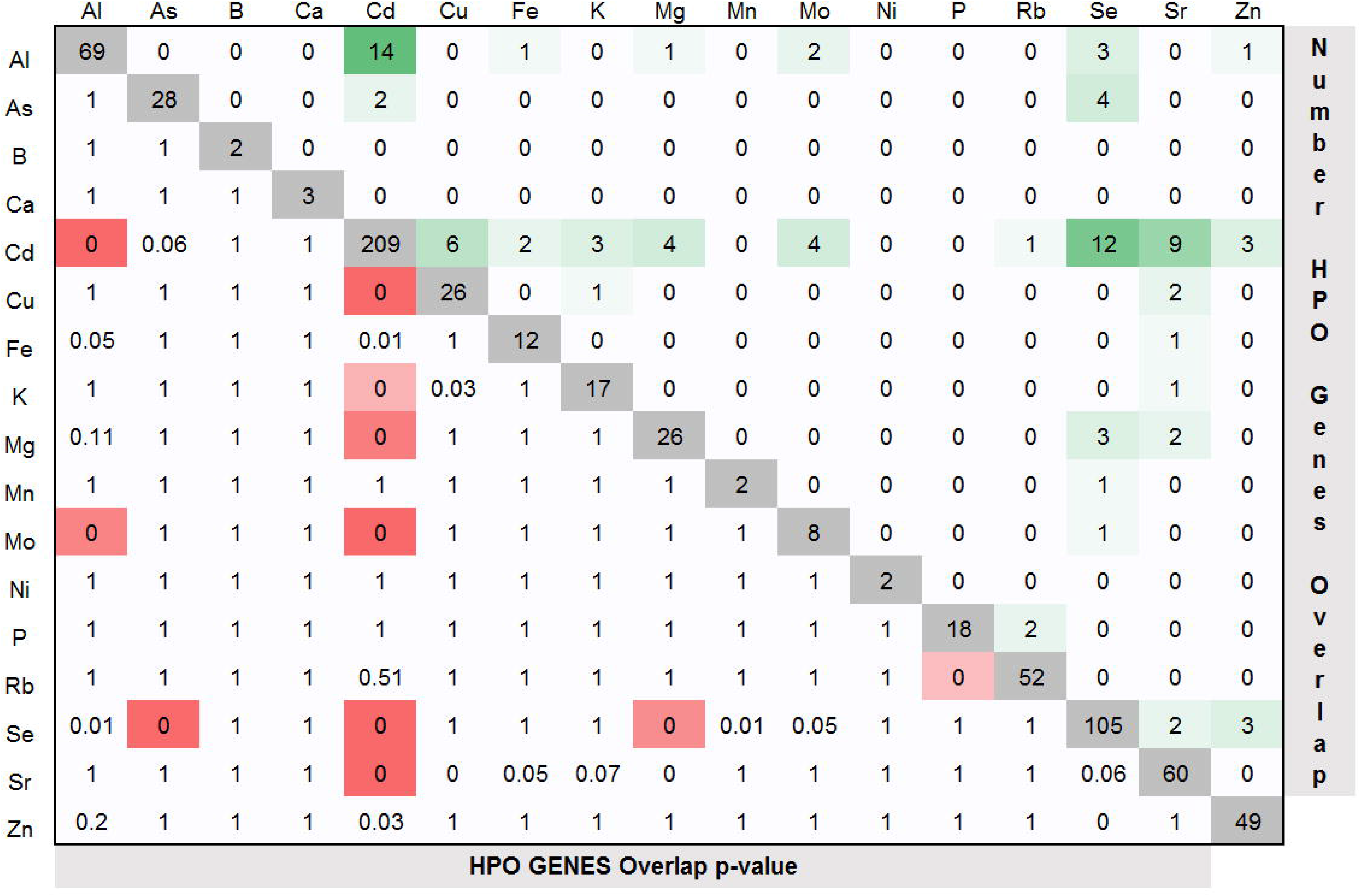
Element HPO candidate gene overlap heatmap. Overlap between the 610 HPO genes discovered between different elements by either density or locality and in any network. The diagonal (grey) shows the number of HPO genes discovered for each element. Values in the upper triangular region show the number of genes that overlap between elements. Cells are shaded green based on the total number of genes they share. The values in the lower triangle designate the *p*-values (hypergeometric) for overlap between the two sets of HPO genes. Red shaded cells indicate significance with Bonferroni correction for multiple testing. Supports Figure 6.

**Supp. Figure 8.**
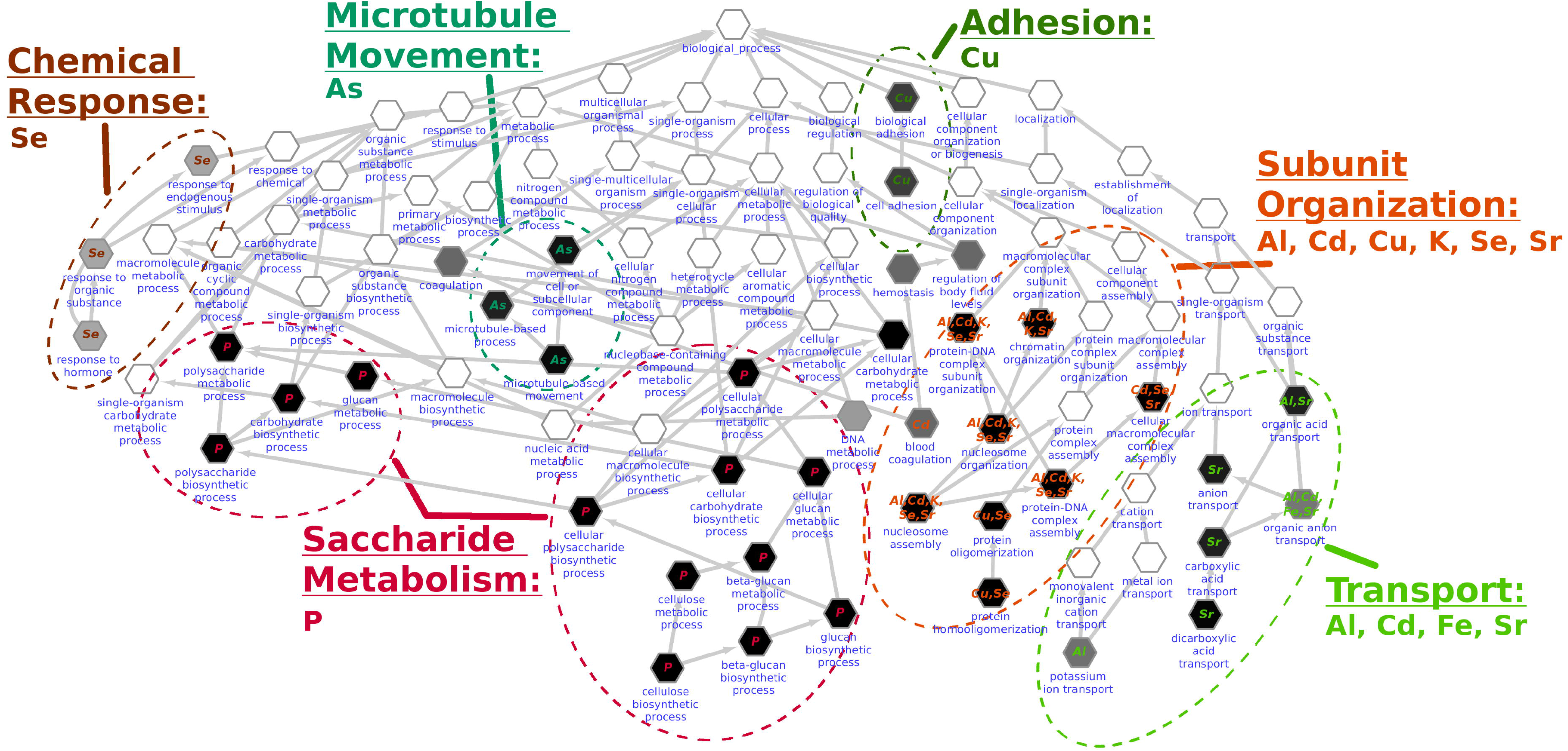
GO biological process enrichment for the ionome. The HPO+ gene sets were analyzed for GO enrichment in the “biological process” namespace. Each node represents a GO term organized hierarchically in a tree with directed edges designating parent terms. Shaded terms were enriched for HPO+ genes (*p* ≤ 0.05; hypergeometric). Dotted ovals represent curated functional terms describing the enriched nodes in different clades of the tree. Each clade is annotated with the ionomic terms that were represented in the GO enrichment. Supports Figure 6.

**Supp. Figure 9.**
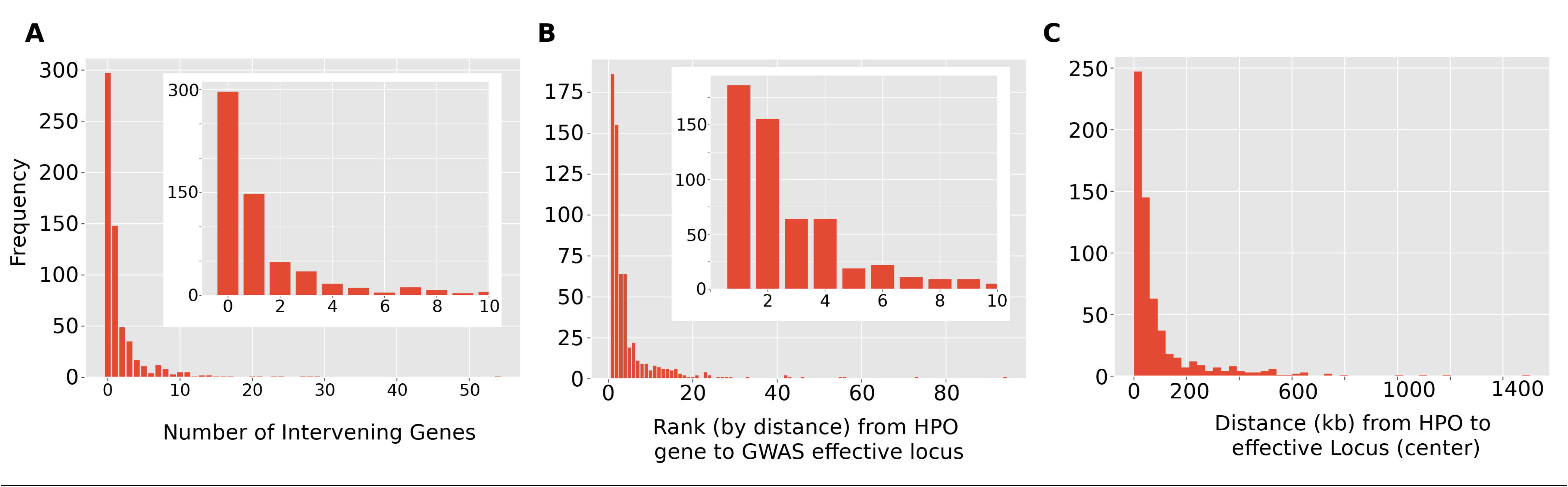
Number of intervening genes between HPO gene and GWAS locus. The distribution of positional candidates and HPO genes. Panel **(A)** shows the distribution in the number of positional candidates between each of the 610 HPO genes and an effective locus (note: intervening gene could also be an HPO gene). Panel **(B)** shows candidate genes near GWAS SNPs, ranked by their absolute distance to effective loci. The distribution shows the rank of the absolute distance (either upstream or downstream) of HPO genes. In both panels, the inset plot shows the lower end of the distributions. Panel (C) shows the distance between the center of HPO genes and the center of the effective locus identified by GWAS. Supports Figure 6.

## Supplementary Text

### Validating density and locality

Density and locality were measured for subnetworks consisting of the set of genes co-annotated to each GO term and compared to scores from 1,000 random sets of genes of the same size (see Table 1; Supp. Table 1 for full data). In total, 818 GO terms of the 1078 tested (76%) were composed of gene sets that were significantly co-expressed (*p* ≤ 0.01) in at least one network using density or locality relative to the randomized gene lists of the same size. Broken down by network as well by Co-expression score, there was substantial Co-expression among GO terms for both density and locality in each network. Density was significant for the most GO terms in the ZmRoot network, while locality performed best in ZmPAN (see Table 1). Considering terms captured by both scores or by either score, overlap between the two Co-expression metrics was comparable. As previously reported (Schaefer et al., 2014), GO terms that exhibit strong Co-expression between members often do so in only a subset of the networks (Supp. Table 1). Thus, both the biological context of the expression data and nature of the Co-expression score influence the subset of GO terms with significantly Co-expression. Overall, while density and locality recover different GO terms, there are substantially more GO terms with significantly co-expressed genes, for either score, than those found by size-matched randomly generated sets of genes (Supp. Table 1).

In addition to detecting strong Co-expression among genes previously annotated by functional processes, unsupervised network clustering using the Markov Cluster algorithm (Dongen, 2000) showed distinct modules within each network (Supp. Table 2). A large number of clusters were significantly enriched for genes that are co-annotated for the same GO term (hypergeometric *p*-value ≤ 0.01; Supp. Table 3). Not all clusters identified previously annotated gene sets. Many strongly co-expressed clusters lacked any previously annotated function (Table 2; Supp. Table 3) potentially identifying novel co-regulated biological processes. Additionally, all networks exhibited a truncated power law distribution in the number of significant interactions (degree) for genes in the network (Supp. Figure 1–3), which is typical of biological networks (Ghazalpour et al., 2006).

### Enrichment analysis of HPO and HPO+ candidate gene sets

GO enrichment was performed among HPO candidate gene lists for each element. Sr was enriched for anion transport (GO:0006820; *p* ≤ 0.008) and metal ion transmembrane transporter activity (GO:0046873; *p* ≤ 0.015). Possibly due to insufficient functional annotation of the maize genome, these enrichment results were limited, and zero elements passed a strict multiple-test correction (Bonferroni). To compensate for the sparsity of annotations, we used the HPO gene set discovered for each trait to identify the set of highly connected Co-expression network neighbors, designated the HPO+ sets. Inclusion in HPO+ was determined by a gene’s aggregate connectedness to the HPO set (see Methods). The HPO+ sets for several of the ionomic traits showed strong GO enrichments, many of which had terms that passed strict multiple-test correction, including Al, As, Cd, Cu, Fe, K, P, Se, Sr, and Zn (Supp. Table 11). Several of the enriched GO terms were common across HPO+ sets for different elements (Supp. Figure 9). For example, we found enrichment for a collection of GO terms related to ion transport (GO:0006811), including anion transport (GO:0006820), potassium ion transport (GO:0006813), and others (GO:0015849, GO:0015711, GO:0046942, GO:0006835), which were supported by enrichments from multiple elements (Al, Cd, Fe, Sr) (see Supp. Figure 9; “Transport” cluster). We also observed a set of six elements whose HPO+ sets (Al, Cd, Cu, K, Se, Sr) were enriched for GO terms related to chromatin organization (e.g., GO:0006325, GO:0071824, GO:0034728, GO:0006334; see Supp. Figure 9, “Subunit Organization” cluster). This may result from changes in cell cycle or endoreduplication control in roots, which is expected to alter the accumulation of multiple elements (Chao et al., 2011).

Several of the observed GO enrichments were trait specific, including collections of GO terms reflecting “chemical response” (Se), “microtubule movement” (As), “adhesion” (Cu), and “saccharide metabolism” (P). For example, the “saccharide metabolism” collection of GO term enrichments was driven by five HPO+ genes for P, one of which was *tgd1* (GRMZM2G044027; see Supp. Table 11). Mutations in the *Arabidopsis thaliana* ortholog of *tgd1* caused the accumulation of triacylglycerols and oligogalactolipids and showed a decreased ability to incorporate phosphatidic acid into galactolipids (Fan et al., 2015), which may alter P accumulation directly or via phosphatidic acid signaling (Katagiri et al., 2005). TGD1 is an ATP-binding cassette (ABC) transporter known to transport multiple substrates, including inorganic and organic cations and anions (Roston et al., 2012). The *tgd1* gene was present in the HPO set, and four other genes were identified as strongly connected neighbors (HPO+) in the Co-expression network. Two genes, GRMZM2G018241 and GRMZM2G030673, are of unknown function, and the other two, GRMZM2G122277 and GRMZM2G177631, are involved in cellulose synthesis. The enriched GO terms demonstrated idiosyncrasies in automated annotation approaches. Terms related to “blood coagulation” and “regulation of body fluid levels” were recovered, which were likely due to annotations translated to maize genes on the basis of sequence homology to human genes. While these term descriptions are not applicable to plant species, the fact that these terms contained HPO genes and exhibited strong network Co-expression suggests that annotations assigned through sequence similarity still capture underlying biological signals for which the assigned name is inappropriate (see Discussion).

In general, using Co-expression networks to expand the neighborhood of the high-confidence candidate causal genes and then assessing the entire set for functional coherence through GO enrichment is a productive strategy for gaining insight into what processes are represented. Yet this approach is particularly challenging in the annotation-sparse maize genome, where only *~1%* of genes have mutant phenotypes (Lawrence et al., 2004). GO terms were too broad or insufficiently described to distinguish causal genes. However, the terms discovered here contain genes that act in previously described pathways known to impact elemental traits. With greater confidence that subnetworks containing HPO genes contained coherent biological information, we refined our analysis by curating HPO genes for their involvement in specific biological processes, namely, those that are known or suspected to affect the transport, storage, and utilization of elements.

### Gene Co-expression analysis of D9

Genes co-expressed with D9 were investigated to determine which were associated with ionomic traits, in particular, seed Cd levels. In the ZmRoot network, D9 was strongly co-expressed with 38 other HPO genes (Figure 7A). Among these were the maize Shortroot paralog (GRMZM2G132794) and a second GRAS domain transcription factor (GRMZM2G079470). Both of these, as well as the presence of many cell-cycle genes among the co-expressed genes and ionomics traits affecting genes, raised the possibility that, like in *Arabidopsis*, DELLA-dependent processes, which are responsive to GA, shape the architecture of the root and the maize ionome. In *Arabidopsis*, DELLA expression disrupts Fe uptake, and loss of DELLA prevents some Fe-deficiency-mediated root growth suppression (Wild et al., 2016). Our finding that constitutive DELLA activity in the roots results in excess Fe, as determined by the *D9-1* and *D8-mpl* mutants, points to a conserved role for the DELLA domain transcription factors and GA signaling for Fe homeostasis in maize, a plant with an entirely different Fe uptake system than *Arabidopsis*. However, the direction of the effect was opposite to that observed in *Arabidopsis*. Future research into the targets of the DELLA proteins in maize will be required to further address these differences.

Remarkably, the HPO Co-expression network associated with D9 in the roots contained three genes with expected roles in the biosynthesis and polymerization of phenylpropanoids (Monaco et al., 2013). The genes encoding enzymes that participate in phenylpropanoid biosynthesis, *ccr1* (GRMZM2G131205), the maize *ligB* paralog (GRMZM2G078500), and a laccase paralog (GRMZM2G336337), were co-expressed with D9. The extradiol ring cleavage dioxygenase encoded by the *ligB* gene (GRMZM2G078500), which from all angiosperms was known to be required for the formation of a pioneer specialized metabolite of no known function in *Arabidopsos*, was linked to QTL for multiple ions including Cd, Mn, Zn, and Ni. The *laccase-12* gene (GRMZM2G336337) was also a multi-ionomic hit with linked SNPs affecting Cd, Fe, and P. The cinamoyl CoA reductase gene, *ccr1* (GRMZM2G131205, was only in the HPO set for Cd. Transcripts co-expressed with D9 also were identified in the ZmPAN network. Consistent with the hypothesis that maize DELLA-domain transcription factors regulate the type II iron uptake mechanism used by grasses, the *nicotianamine synthase3* gene (GRMZM2G439195, ZmPAN-Cd), which is required for making the type II iron chelators, was both a Cd GWAS hit and substantially co-expressed with D9 in the ZmPAN network, such that it contributed to the identification of d9 as an HPO gene for Cd.

### Previously described HPO genes and their effects on the ionome

We expect that changes to seed compartment proportions or the production of major storage constituents will alter seed ionomic content. Within the NAM population, functional variation for *su1* can be found in the B73 x IL14H subpopulation. For this reason, six IL14H recombinant inbred lines (RILs) that were still segregating for the recessive *su1* allele were previously tested for ionomic effects (Baxter et al., 2014). This demonstrated that segregation for a loss of function allele at *su1*, on the cob, affected the levels of P, S, K, Ca, Mn, Fe, As, Se, and Rb in the seed (Baxter et al., 2014). Previous analysis of lines segregating *su1* allele in the IL14H RIL population and measured in the NAM panel, four were associated with *su1* variation in the association panel. It is possible that *su1*, which is expressed in multiple plant compartments including the roots, might also have effects throughout the seed ionome beyond a dramatic loss of seed starch. This may result from coordinate regulation of the encoded isoamylase and other root-expressed determinants of S and Se metabolism, or from unexpected coordination between root and seed expression networks. The finding that HPO network neighbors for P were enriched for carbohydrate biosynthetic enzymes favors the former of these two hypotheses (see Supp. Figure 9).

Our combined analysis of loci-linked GWAS SNPs and gene Co-expression networks identified a large number of HPO genes associated with Se accumulation. Several genes with known demonstrated effects on the ionome, or known to be impacted by the ionome, were identified within this HPO set. For example, GRMZM2G327406, encodes an adenylyl-sulfate kinase (*adenosine-5′-phosphosulfate [APS] kinase 3*), which is a key component of the sulfur and selenium assimilation pathway and plays a role in the formation of the substrate for protein and metabolite sulfation (ZmRoot-Se). At another locus, Camoco identified a cysteine desulfurase (GRMZM2G581155), critical for the metabolism of sulfur amino acids and the biosynthesis of the 21st amino acid selenocysteine, as an HPO gene (ZmRoot-Se).

Based on the work of Chao et al. in *Arabidopsis*, alterations in cell size and cell division in the root are expected to have effects on K accumulation in leaves (Chao et al., 2011). Two of the four subunits of the polycomb repressive complex 2 (PRC2), known to act on the cell cycle via the retinoblastoma-related proteins (RBRs), were identified as HPO genes for the K analog Rb. Both *msi1* (GRMZM2G090217; ZmSAM-Rb) and *fie2* (GRMZM2G148924; ZmSAM-Rb), members of the Polycomb Repressive Complex2, are co-expressed in the ZmSAM network. The RBR-binding E2F-like transcription factor encoded by GRMZM2G361659 (ZmSAM-Rb) was also found, a further indication that cell-cycle regulation via these proteins’ interactions could provide a common mechanism for these associations. Histone deacetylases from the RPD3 family are known to interact with RBR proteins as well. The RPD3-like *histone deacetylase 2* gene from maize was identified in the same HPO set (GRMZM2G136067; ZmSAM-Rb). The *Arabidopsis* homologs of both *msi1* and *histone deacetylase2* have known roles as histone chaperones, and the latter directly binds histone H2B. Remarkably, *histone H2B* (GRMZM2G401147; ZmSAM-Rb) was also an HPO hit. Lastly, the actin-utilizing-SNF2-like *chromatin regulator18* gene (GRMZM2G126774; ZmSAM-Rb) was identified as yet another SAM-Rb hit. This mirrors the similar finding of GO enrichment for chromatin regulatory categories in the HPO+ enrichment analysis presented above. Taken together, these demonstrate a strong enrichment for known protein-protein interactors important for chromatin regulation and cell-cycle control among the HPO set for the K analog Rb.

Several annotated transporters were identified in the HPO sets for multiple elements: a putative sulfate transporter (GRMZM2G444801; ZmRoot-K), a cationic amino acid transporter (AC207755.3_FG005; ZmPAN-Cd, ZmPAN-Mo), and an inositol transporter (GRMZM2G142063; ZmRoot-Fe, ZmRoot-Cd, ZmRoot-Sr).

Cadmium is well measured by ICP-MS and affected by substantial genetic variance (Ziegler et al., 2017). We detected the largest number of HPO candidate genes for Cd (209 genes; see Figure 6). Among these were the maize *glossy2* gene (GRMZM2G098239; ZmPAN-Cd), which is responsible for a step in the biosynthesis of hydrophobic barriers (Tacke et al., 1995). This implicates the biosynthesis and deposition of hydrophobic molecules in accumulation of ions and may point to root processes, rather than epicuticular waxes deposition, as the primary mode by which these genes may affect water dynamics. An ARR1-like gene, GRMZM2G067702, was also an HPO gene associated with Cd (ZmRoot). Previous work has shown that ARR genes from *Arabidopsis* are expressed in the stele, where they regulate the activity of HKT1 (Mason et al., 2010). This gene was expressed at the highest level in the stele at 3 days after sowing.

